# Calves function as social magnets in a species with high fission-fusion dynamics, the Asian elephant

**DOI:** 10.64898/2026.11.01.763273

**Authors:** T. Revathe, S. Nandini, P. Keerthipriya, Hansraj Gautam, Ankana Sanyal, Chowdari Jabili, Divya Choudhary, S. Anvitha, T.K. Athira, T.N.C. Vidya

## Abstract

Individuals living in fission-fusion societies may face unique challenges in maintaining social relationships, as the repeated fission of social units can limit opportunities for social interaction. However, the presence of young offspring may often necessitate cooperation among individuals. To understand whether offspring promote sociality in fission-fusion species, we studied wild Asian elephants (*Elephas maximus*) in southern India. Using nearly 5900 female group sightings collected over 9 years, we tested whether calf presence predicted female group size and associations. The numbers of adult, young adult, and subadult females in groups were 2-2.5-fold higher in calf presence than absence, which may facilitate collective defence of calves. Furthermore, the number of females without their own dependent offspring, who often provide allomaternal care, was also higher in calf presence than absence. Social network analysis revealed that adult females formed more numerous and stronger associations in calf presence than absence, which may facilitate consistent allomaternal care. Our results thus suggest that the need to protect calves and to provide allomaternal care likely promote sociality among female elephants. We propose that dependent offspring may function as social ties in fission-fusion species, contingent upon the prevailing conditions and the socioecological pressures that have shaped their social systems.

## 1. Introduction

Group living is widespread across many mammals and can confer several benefits including increased access to resources, improved cooperation, and reduced risk of predation [1,2]. However, group living also entails costs such as increased resource competition, disease transmission, and infanticide risk [1,2]. Fission-fusion dynamics is thought to provide a means of flexibly balancing these competing costs and benefits [3–5]. Individuals in fission-fusion societies flexibly adjust their group size and/or composition (i.e., social organisation [1,6]) in response to spatiotemporal variation in ecological conditions [7–16] through the splitting into and merging of subgroups [3–5]. By causing repeated, and extended, separations among group members, fission-fusion dynamics can present unique challenges for individuals in terms of opportunities to interact and, thereby, in maintaining social relationships [5]. Therefore, identifying the factors that facilitate the persistence, and strengthening, of social relationships, despite separations, is crucial for understanding sociality in fission-fusion societies. Such studies may also provide insights into the conditions that favour the evolution of fission-fusion societies.

The presence of dependent offspring is one of the factors that has the potential to repeatedly bring together individuals, particularly females, in mammalian fission-fusion societies. The successful development and survival of dependents to reproductive age requires nutritional input, locomotory support, protection against predation or infanticide, social support, and opportunities for social learning and socialisation during the developmental period [17–19]. In species in which primarily the mother takes care of her dependent offspring, other females in the group may fulfil roles that cannot be achieved solely by the mother, such as assistance during birth [20] or collective defence of the dependent [21,22]. In species with allomothering, where females other than the biological mother provide care to conspecific young [23], mothers and allomothers likely have to repeatedly associate to facilitate consistent allomaternal care, which may strengthen existing relationships [24–26]. Therefore, the need for cooperation among females in the presence of dependent offspring may promote female gregariousness (i.e., the tendency to associate with others, which can resultantly shape group size) and the maintenance of preferential social relationships. Thus, dependent offspring may have the potential to shape female social structure (i.e., the patterning of social relationships [6]).

To this day, the effect of the presence of dependent offspring on two main aspects of female sociality (i.e., social organisation and structure [5]) remains largely overlooked. The few studies examining group size [7–9,11,12,27–29] have yielded contrasting results. In spotted hyena (*Crocuta Crocuta*), for example, females with small cubs avoided conspecifics, likely to reduce the risk of adult female-driven infanticide [7]. In giraffe (*Giraffa tippelskirchi*), females adopted a calf-hiding strategy when predator density was high, and thus group sizes decreased with an increase in the proportion of calves in the group, specifically in habitats with dense vegetation [11]. In contrast, studies on cetaceans and a few other species found that groups with dependent offspring were larger than those without them [8,9,12,27–29]. However, in the latter group of studies it is difficult to ascertain whether the presence of dependent young promoted female gregariousness, as confounds such as season were not controlled for and/or the definition of group size also included the dependent offspring themselves and even males who do not provide care in these species.

With respect to female social structure, a study on sperm whales (*Physeter macrocephalus*) found that several social network metrics among females increased with the birth of a calf in the social unit [25]. With the analysis being limited to a single group of five females, it is unclear how generalisable these results are at the population level, although the reported raw values across six social units [25, table 3] suggest that calves may increase sociality among female sperm whales. Another study on the plurally breeding degus (*Octodon degus*) with communal care found that association strength among females was higher during the lactation than the mating period [30]. However, increased association strength was not found to benefit females in terms of per capita pup production [30]. Thus, these two studies provide only limited, yet valuable, evidence that the need for allomaternal care may shape female sociality in fission-fusion species.

To address the dearth of studies clearly determining the effect of dependents on female sociality, and to identify potential reason(s) behind any such effect, we investigated whether and how female social organisation and structure differ between the presence and absence of calves (<1 year of age) using the Asian elephant (*Elephas maximus*) as the study system. Below, we detail the reasons why Asian elephants are an excellent model system to address this question.

Female Asian elephants are organised into distinct clans or communities that display a high degree of fission-fusion dynamics [15,31]. The clan is the most inclusive social unit and is composed of philopatric females and their dependent offspring and subadults of both sexes [31]. Clans are likely stable in adult female membership over time, unless (very rarely) they dissolve due to the death of most or all clan members [32], and females are often, although not always, related within clans [33,34]. Fission-fusion dynamics occur within clans, with females of a clan being seen in the form of groups (see Methods) that are not stable in composition [31]. Thus, a group is usually a subset of the clan, and rarely the entire clan, and group size and composition change over timescales of hours to days [15,31]. Overall, females maintain extensive associations, with variable numbers of weak and strong social ties within their clans [14,15,31,35]. Weak dominance relationships have been observed between females within clans [36], whereas intense agonistic interactions are found between clans [37]. In contrast to females, males disperse from their natal clans when they are about 10 years old, and thereafter form only temporary associations with females [38,39]. Males do not provide offspring care in elephants [40]. There are no known cases of infanticide by either sex [41].

Calves are able to follow their mothers soon after birth [42], but have a lengthy developmental period, spanning at least 2-3 years [42,43], and they are vulnerable to tiger attacks [44–46]. Calves are thus heavily dependent on their mothers and allomothers for nutrition (consumption of milk almost always only from mothers; tolerated taking of solid food from both mothers and allomothers [24]), protection against predation, social support, and likely also for social learning [24,26,42,43,46–48]. The need for calf protection and alloparental care may thus promote sociality among Asian elephant females [47]. To test this hypothesis, we used a decade of group size and association data collected on the long-term studied Kabini elephant population in southern India [32,49,50] and investigated three specific questions.

### (a) Q1: Does calf presence predict group size?

We did not have an *a priori* prediction for the effect of calves on the number of females in a group (i.e., female group size). Due to potential constraints on group size in the study population [31,51], it was possible that female group size would not increase in calf presence. However, it was also possible that female group size would temporarily increase in the presence of calves, despite average group sizes being small overall, due to anti-predatory benefits in a habitat with high tiger density [52]. In such a case, the number of adult females of any age-class might be expected to increase in calf presence, as dependent offspring are usually collectively defended by females [46,47]. An increase in female group size might also occur through the addition of primarily younger females (young adults or subadults; see Methods), specifically those without their own calf or young juvenile (1-<2 years of age), as such females often extend allomaternal care in the study population [24,26]. We predicted that the number of adult and subadult males associating with female groups would be similar between groups with and without a calf, as there is neither infanticide nor care by males.

### (b) Q2: Does calf presence predict group size experienced by individual adult females?

In populations with a high variance in group sizes, a majority of the individuals will experience group sizes larger than the average, and hence must be weighted proportionately. The group size that an average individual experiences would then be a better measure of animal grouping tendencies than average group size [53]. As female group sizes in the study population are highly variable [31], in addition to examining the number of females in a group, we examined whether the number of females experienced by individual adult females in a group (i.e., experienced group size) differed between the presence and absence of calves.

### (c) Q3: Are adult female association patterns different between calf absence and presence?

As allomaternal care is important for offspring at least during the first year of life [24,46,47], we predicted that adult females would increase their strengths of associations with certain females in the presence of calves. An increase in the number of associates might also be expected if the group size increased in the presence of calves. However, even in the absence of group size changes, individual-based fission-fusion dynamics [31] would allow for certain females to take the place of others, possibly resulting in differences in association strengths between calf presence and absence. We tested two other social network metrics (see Methods) and did not have *a priori* predictions for them.

## 2. Methods

### (a) Study site and field data collection

To investigate the effect of calves on female sociality, we studied the Kabini Asian elephant population [32,49], which inhabits the Nagarahole and Bandipur National Parks and Tiger Reserves (currently Nagarahole Tiger Reserve: 11°45 –12°15 N, 76°05 –76°25 E, 643.35 km^2^ core area; currently Bandipur Tiger Reserve: 11°35 –11°55 N, 76°12 –76°51 E, 872.24 km^2^ core area) in southern India. The parks mainly comprise dry and moist deciduous forests, and are separated by the Kabini Reservoir on the River Kabini. The parks experience a dry (mid-December to mid-June) and a wet season [14], with the receding backwaters of the reservoir forming an open grassland during the dry season, which attracts large numbers of elephants and other herbivores. The Kabini elephant population has been studied since 2009 [32,49], and over 1400 elephants have been individually identified as part of the Kabini Elephant Project [54]. For this work, we used the female group sighting data (henceforth, sightings) collected from March 2009 to June 2018. Data were collected between about 6:15 AM and 5:45-6:45 PM (depending on daylight hours and field permits). We sampled fixed routes in the backwaters and forests in a stratified manner, see [31,49] for more details.

Elephants were individually identified based on a combination of natural physical characteristics such as ear shape and length, nicks, holes, and tears on ears, back shape, tail length and tail hair brush shape, and, in the case of males, tusk length, shape, and symmetry [54]. Individuals were aged based on their shoulder height, body length, and skull size, among other measures [54], with estimated errors between 5 and 10 years (usually) for adults, see [50,54]. We defined female elephant groups as a set of one or more adult females (at least 10 years old) within 50-100 m of one another that showed coordinated movement and behaviours [31]. Adult females could be accompanied by subadults (subadult females: 5–<10 years of age, subadult males: 5–<15 years of age), juveniles (1–<5 years of age), and calves (<1 year of age) of both sexes, and sometimes by adult males (≥15 years old). Rarely, one or more subadult females formed female groups without any adult female, and could be accompanied by individuals of younger age-sex classes. Following the ‘gambit of the group’ approach, we considered all the individuals in a group to be associating with one another. We considered adult and subadult males to associate with a female group if they fed within about 10 m of the group or interacted with the group [39].

We identified clans through modularity-based community detection (Louvain method) [55] in association networks [31]. Female groups almost always consisted of females of a single clan [31]. However, over the years, because of the near-extinction of a few clans or other reasons, the surviving females joined other clans [32, Kabini Elephant Project, unpublished data], although they may not have been fully integrated into the new clans. However, as between-clan associations were rare and between-clan interactions were generally agonistic in the study population [34,37], only rarely would calves be expected to have an (enduring) positive effect on associations among females between clans. Therefore, for the purposes of this study, we used only single-clan sightings (i.e., sightings in which females of a single clan associated with one another, with clan as defined over the first half of the study, see [31]), as we were interested in determining the effect of calves on within-clan female sociality.

### (b) Data analysis

To quantify the number of females and males in a group, the number of females experienced by adult females, and adult female association, we used independent sightings in which all the individuals were age-sex classified. Independent sightings were those that were separated by at least 2.5 hours because at this timepoint, the probability of a female group changing in composition or not was roughly similar [31].

We fit Generalized Linear Mixed Effects Models (GLMM) to the number of females and males in a group and the number of females experienced by adult females. As these were count data, if there was no overdispersion, we used a Poisson error distribution. Otherwise, we used a negative binomial error distribution. When the response did not include zero, we used a truncated Poisson distribution. We used the log link for all the models. All these analyses were run in R (version 4.6.0) [56] using the package ‘glmmTMB’[57]. The group size and experienced group size model structures are provided along with the model estimates in the supplementary materials (referenced in Results).

#### (i) Q1: Does calf presence predict group size?

To investigate whether the number of females and males differed between groups with and without calves, for each sighting, we calculated the number of 1) adult females, 2) young adult females (≥10 to <15 years), and 3) subadult females (≥5 to <10 years), which were our dependent variables of interest (models: m_AF, m_YAF, and m_SAF, respectively). To more directly test whether the need for allomaternal care increases female sociality, we quantified the number of adult and young adult females without their own dependent offspring (<2 years of age) in each sighting (models: m_AF1 and m_YAF1, respectively), as only females without dependent offspring generally provide allomaternal care in Asian elephants [24,26,47]. As subadult females would generally be too young to give birth [50], we did not repeat the above analysis for females of this age-class. We used calf status in each sighting as a categorical predictor (absence/presence). Calf presence means that at least one calf was present in the group. Additionally, as the number of adult, young adult, or subadult females that could join a group would depend on the respective numbers of such females available in the clan (supplementary material, S1, table S1), clan demography had to be controlled for in the models. We did so by including the number of females of a specific age-class (adult/young adult/subadult, depending on the model) in the clan as a control predictor (discrete counts).

If female group size increases to protect calves against predation or to provide allomaternal care, we should not expect to see a systematic increase in the number of males in a group in calf presence. To test this, for each sighting, we calculated the number of males that were at least 10 years old (who would have, therefore, begun the process of dispersal and would be able to associate with different female groups) and tested whether these numbers were different between calf presence and absence (model: m_M).

In all the models with the number of females as a dependent variable, we included a random intercept of clan. In order to reliably estimate the random effect of clan, we used the sightings of only those clans that were sighted at least five times each year in both calf presence and absence (note that not all clans were sighted each year; supplementary material, S2, table S1). We also included a random intercept of year in all the models. We included all theoretically identifiable random slopes so as to keep the type-I error rate at the nominal level of 5% [58,59]. Seasonal differences in group size and association measures were not found in the Kabini Asian elephant population after accounting for clan identity, despite strong seasonality in habitat and space use [14]. Therefore, we did not control for season in our models.

#### (ii) Q2: Does calf presence predict group size experienced by individual adult females?

To test the group size that an individual experiences, we calculated the group size experienced by each adult female in each sighting as: 1) the number of adult females (i.e., including the focal adult female), 2) the number of young adult females (i.e., including the focal adult female if she was ≥10 to <15 years), and 3) the number of subadult females (models: mE_AF, mE_YAF, and mE_SAF, respectively). Furthermore, as we were interested in testing whether allomothering is a reason for any difference in experienced group size, we also calculated the 4) the number of other adult females (i.e., excluding the focal adult female) without their own dependent offspring and 5) the number of other young adult females (i.e., excluding the focal young adult female) without their own dependent offspring (models: mE_AF1 and mE_YAF1, respectively) that an adult female experiences in a sighting. Calf presence/absence here refers to the status in the sighting, irrespective of whose calf it was. Since calf status was not a significant predictor of the number of males in a group (see Results), we did not test whether calf status predicted the number of males experienced by adult females.

Adult females were repeatedly sampled within and across years, and the experienced number of females was quantified for each adult female in a given sighting. Therefore, to avoid pseudo-replication, in addition to the predictors and random intercepts described above for the group size analysis, we included random intercepts of adult female identity and sighting ID. As before, we built models with maximal random effects structure. As each female belongs to only one clan, we nested female ID within clan. To reliably estimate the random effect of this nested random effect, we included only those clans for which we had data on at least five adult females (over the entire study period) who were sighted at least five times in both calf absence and presence within each year (note that not all females were sighted each year; supplementary material, S2, table S2).

#### (iii) Model checks and data visualisation

When the estimated correlation between random slopes and intercepts was close to the boundary (−1 or +1), indicating unidentifiable correlation parameter, we removed it from our model. To ease model convergence and to get easily interpretable estimates [60], we *z*-transformed all the continuous predictors to a mean of 0 and a standard deviation of 1. After fitting each model, we checked for overdispersion and zero-inflation (none of the models were zero-inflated) using the ‘DHARMa’ package (version 0.5.0) [61], and collinearity (using *vif* function of the ‘car’ package, version 3.1.5; maximum collinearity was 1.01). Following model checks, as an overall test of the effect of fixed effects and to avoid cryptic multiple testing, we compared the full model to the null model (without fixed effects but with the same random effects structure as the full model) using Likelihood Ratio Tests (LRT), as implemented in the *anova* function in R. Only if the full model was significantly better than the null model, we interpreted the model estimates. To test the significance of the predictor, we used the *drop1* function in R, which drops one predictor at a time and compares it to the full model using LRT. We considered the estimate to be significant at *P*<0.05. As a measure of effect size, we report the exponent of the estimate (back-transformed to the original response scale from the log link space). We also report marginal coefficients of determination (i.e., the variance explained by the fixed effects; *R*^2^m) and conditional coefficients of determination (i.e., the variance explained by the entire model; *R*^2^c) estimated using the *r2_nakagawa* function of the ‘performance’ package (version 0.17.0). All the graphs were plotted in R using the ‘ggplot2’ package (version 4.0.3), and the model predictions and their confidence intervals for the graphs were obtained using the *ggpredict* function of the ‘ggeffects’ package (version 2.3.2).

#### (iv) Q3: Are adult female association patterns different between calf absence and presence?

For the association analyses, we used only adult females because subadult female associations might not have been independent of those of adult females. We calculated association indices between pairs of adult females separately for sightings with and without calves. Since we were interested in testing differences in within-clan female association patterns between calf presence and absence, we only used those clans for which we had data on at least two adult females. The females used for the analysis were seen at least five times each in calf absence and calf presence (not necessarily their own calf) across the entire study period (supplementary material, S2, table S3). As we did not have data on enough clans to test changes in female association between calf presence and absence within each sampling year (supplementary material, S3, figure S1), the female association analysis could not be carried out for each year. However, note that we were interested in testing the overall effect of calf absence/presence on female social structure in the study population rather than the year-to-year stability in the effect of this predictor.

Association index (AI) was calculated as the simple ratio index, which is the ratio of the number of times two individuals were seen together to the total number of times they were seen [62]. To test whether the distribution of AI values differed between calf absence and presence, we calculated the kurtosis of AI for each condition. Kurtosis is a measure of the tailedness of a distribution – i.e., the frequency of extreme values (high or low) in relation to the spread of the distribution. It is calculated as the fourth central moment (i.e., the average of deviations from the mean raised to the fourth power) divided by the standard deviation raised to the fourth power. It is often interpreted in comparison with a normal distribution, which has a raw kurtosis of 3. A raw kurtosis of >3 means that a distribution has heavy tails and extreme values occur more often than in a normal distribution. We further calculated two network metrics – degree and clustering coefficient – and tested whether they were different between calf presence and absence using sampled randomisation tests. Degree is the number of direct associates of an individual; and clustering coefficient measures the proportion of a focal individual’s direct associates who are also direct associates of one another [63].

AI, kurtosis of AI, and network metrics were each compared between calf presence and calf absence using sampled randomisation tests [64]. We performed 10,000 permutations of the data by randomly marking sightings as either calf present (permuted ‘calf-present’ dataset) or calf absent (permuted ‘calf-absent’ dataset), while preserving the original number of sightings under each category. For each metric, we calculated the observed difference in the metric between the two calf statuses by subtracting the smaller value from the larger value. The differences in the metric between the permuted calf presence and absence datasets were then calculated using the same subtraction direction as for the observed difference. We calculated *P* values as the proportion of differences from the randomised datasets that were at least as extreme as the observed. These calculations and the randomisation tests were carried out using MATLAB R2015b (The MathWorks, Inc, 1984-2015, www.mathworks.com). Association networks of females in the presence and absence of calves were constructed in Gephi 0.11.2 [65].

## 3. Results

### (a) Q1: Groups with calves have a greater number of females but not males

We used a total of 3968 sightings (*N*=1413 with a calf) collected across 10 years on 14 clans (that were sighted at least five times each in calf presence and absence each year) for the group size analysis (supplementary material, S2, table S1). Overall, the full model with all the predictors was significant as compared to the null model for the number of adult females (m_AF: χ2=25.622, *df*=2, *P*<0.001), young adult females (m_YAF: χ2=34.692, *df*=2, *P*<0.001), and subadult females (m_SAF: χ2 = 22.929, *df*=2, *P*<0.001) in a group. We found that groups with a calf had 1.96, 2.21, and 2.52 times as many adult, young adult, and subadult females, respectively, as groups without a calf (supplementary material, S4, table S1-3, Figure 1A-C). We then tested for the number of females in a group without their own dependent offspring, and the full models were again significantly better than the null models for adult females (m_AF1: χ2=9.671, *df*=2, *P*=0.008) and young adult females (m_YAF1: χ2=22.395, *df*=2, *P*<0.001). As before, the numbers of both adult and young adult females without their own dependent offspring were significantly higher in calf-present than calf-absent groups (supplementary material, S4, table S4-5). However, the magnitude of increase of these adult females was considerably lower (1.43 times as many adult females without a dependent offspring of their own in calf-present than calf-absent groups, supplementary material, S4, figure S1) than that of all adult females (as estimated in m_AF). For young adult females also, the magnitude of increase was lower (1.84 times as many young adult females without their own dependent offspring in calf-present than calf-absent groups, supplementary material, S4, figure S1) than that estimated in m_YAF. The full model for the number of adult males in the group was not significantly better than the null model (m_M: χ2=0.034, *df*=1, *P*=0.855). We therefore do not interpret the estimates of this model.

**Figure 1.**
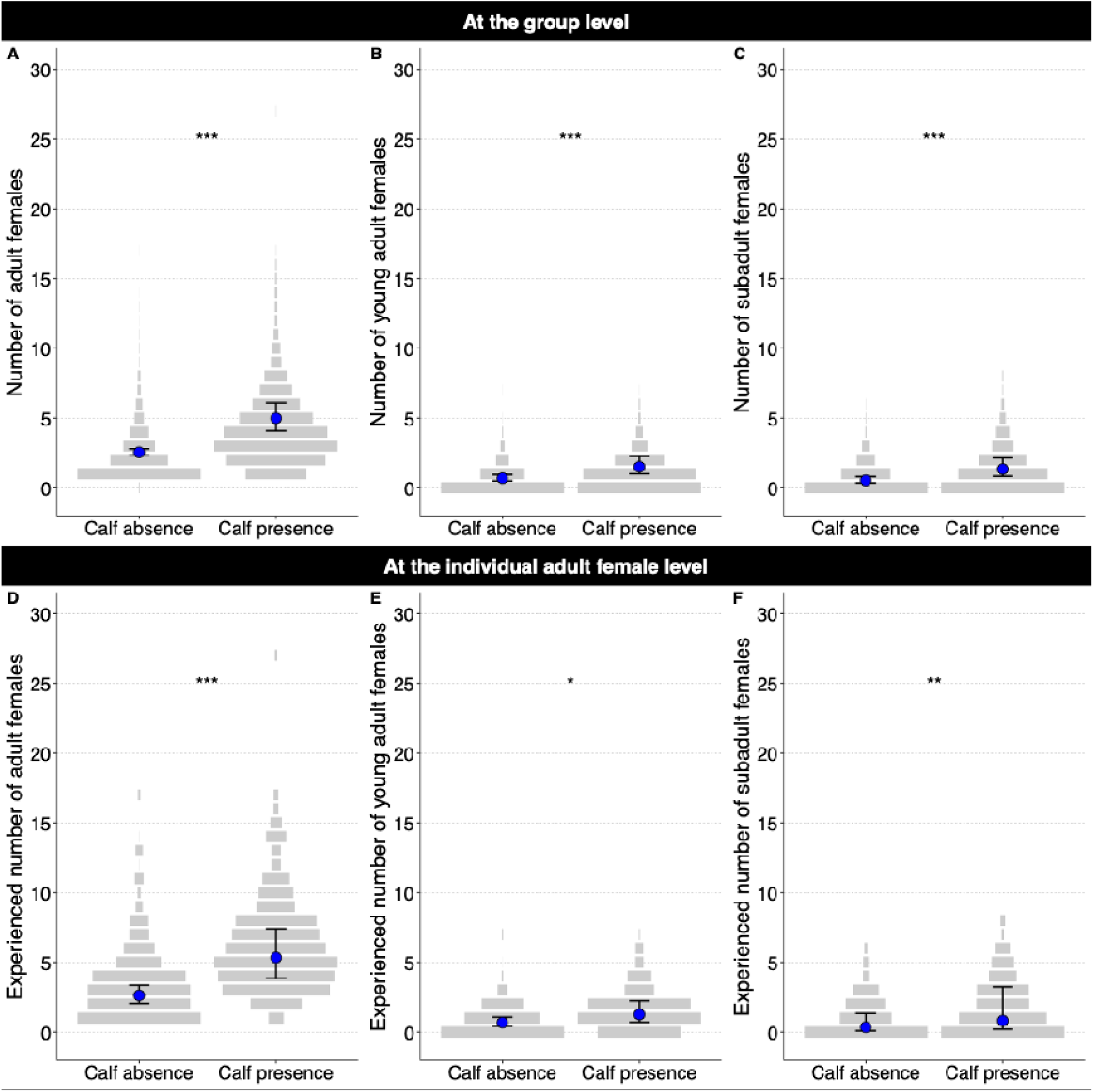
Effect of calves on female group and experienced group sizes. Mean model predictions (blue dot) and the confidence intervals (black bars) are shown for the numbers of (A) adult, (B) young adult, and (C) subadult females in a group and the numbers of these age-class females experienced by adult females, (D), (E), and (F) respectively. The observed data are shown as discrete violin plots (grey bars). The widths of the bars show the relative frequency of observations within each calf status, and width of the widest bar is scaled to be the same between calf presence and absence. Significance codes: <0.001 ‘***’ 0.001 ‘**’ 0.01 ‘*’ 0.05 ‘.’.

### (b) Q2: Group size experienced by individual adult females were higher in calf presence than absence

We used 1946 group sightings of 84 adult females (*N*=867 adult female group sightings in calf presence) from 6 different clans. These females were sighted a total of 4421 times between 2009 and 2018 (excluding 2010; supplementary material, S2, table S2). The full models with all the fixed effects were significantly better than the null models for the experienced numbers of adult females (mE_AF: χ2=14.749, *df*=2, *P*=0.001), young adult females (mE_YAF: χ2=14.244, *df*=2, *P*=0.001), and subadult females (mE_SAF: χ2=7.267, *df*=2, *P*=0.027). Similar to the pattern observed for female group size, we found that calf status had a significant effect on all three experienced group size variables (supplementary material, S5, table S1-3). Specifically, the experienced numbers of adult, young adult, and subadult females in calf presence were 2.04, 1.85, and 2.33 times those in calf absence (Figure 1D-F), respectively.

When we tested the number of other adult or young adult females without their own dependents that an adult female experienced in calf absence and presence, the full models performed significantly better than the null models (mE_AF1: χ2=12.763, *df*=2, *P*=0.002; mE_YAF1: χ2=14.565, *df*=2, *P*=0.001), with calf status being a significant predictor (supplementary material, S5, table S4-5). The experienced numbers of adult and young adult females in calf presence were 1.89 times (lower than that estimated in mE_AF) and 1.97 times (higher than that estimated in mE_YAF) those in calf absence, respectively (supplementary material, S5, figure S1).

### (c) Q3: Adult female associations were stronger in calf presence than calf absence

We quantified AI and social network metrics using a total of 5887 sightings (*N*=1544 sightings in calf presence) of 171 adult females from 17 clans (supplementary material, S2, table S3). AI distributions in the presence and absence of calves were highly skewed (Figure 2A,B). The percentage of non-zero AIs among adult females within clans (each female seen at least 5 times in both calf status) was higher in calf presence (average (SD)=86.27 (18.58)) than in calf absence (77.91 (23.83)), meaning that a greater number of associations among females were realised in calf presence than absence (Figure 2C,D). Furthermore, we found that the average AI between adult females was significantly higher in calf presence (0.013, SD=0.0694) than in calf absence (0.007, SD=0.0425) and the kurtosis of AI was significantly lower in calf presence (71.68) than in calf absence (108.07), based on sampled randomization tests (supplementary material, S6, table S1; Figure 3A,B). There was also a higher percentage of higher AI values within clans in calf presence (average percentage of AI values >0.5 (SD)=22.47 (23.53); Figure 2B,D) than absence (11.24 (24.71); Figure 2A,C).

**Figure 2.**
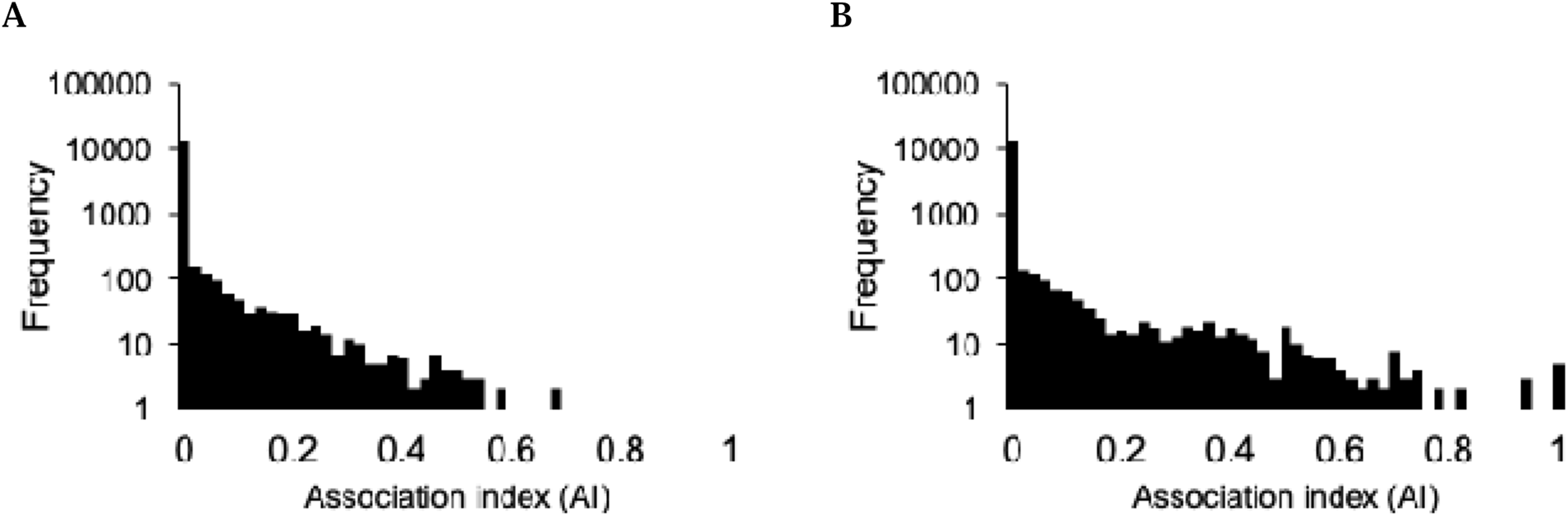
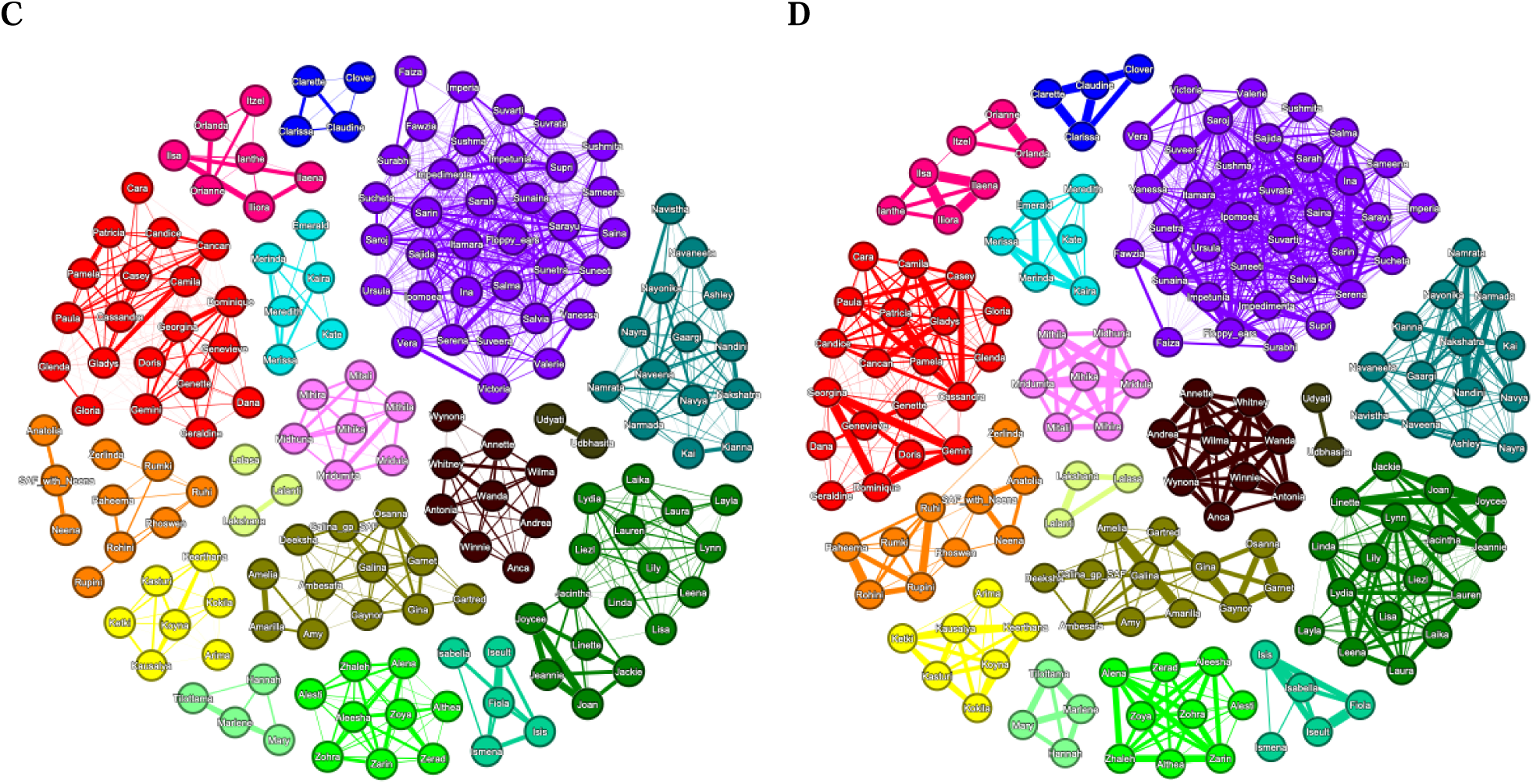
Association indices (AI) and networks of 171 adult females in the absence and presence of calves. Observed AI distributions in (A) calf absence and (B) calf presence are shown. Social networks in (C) calf absence and (D) presence were constructed using the Fruchterman Reingold layout [66]. Each node (i.e., circle) represents an adult female, and the colours represent distinct clans, initially detected as modularity classes. The edges (i.e., lines) between each pair of nodes represent non-zero AI and are weighted by AI values (representing the thickness of the edges).

**Figure 3.**
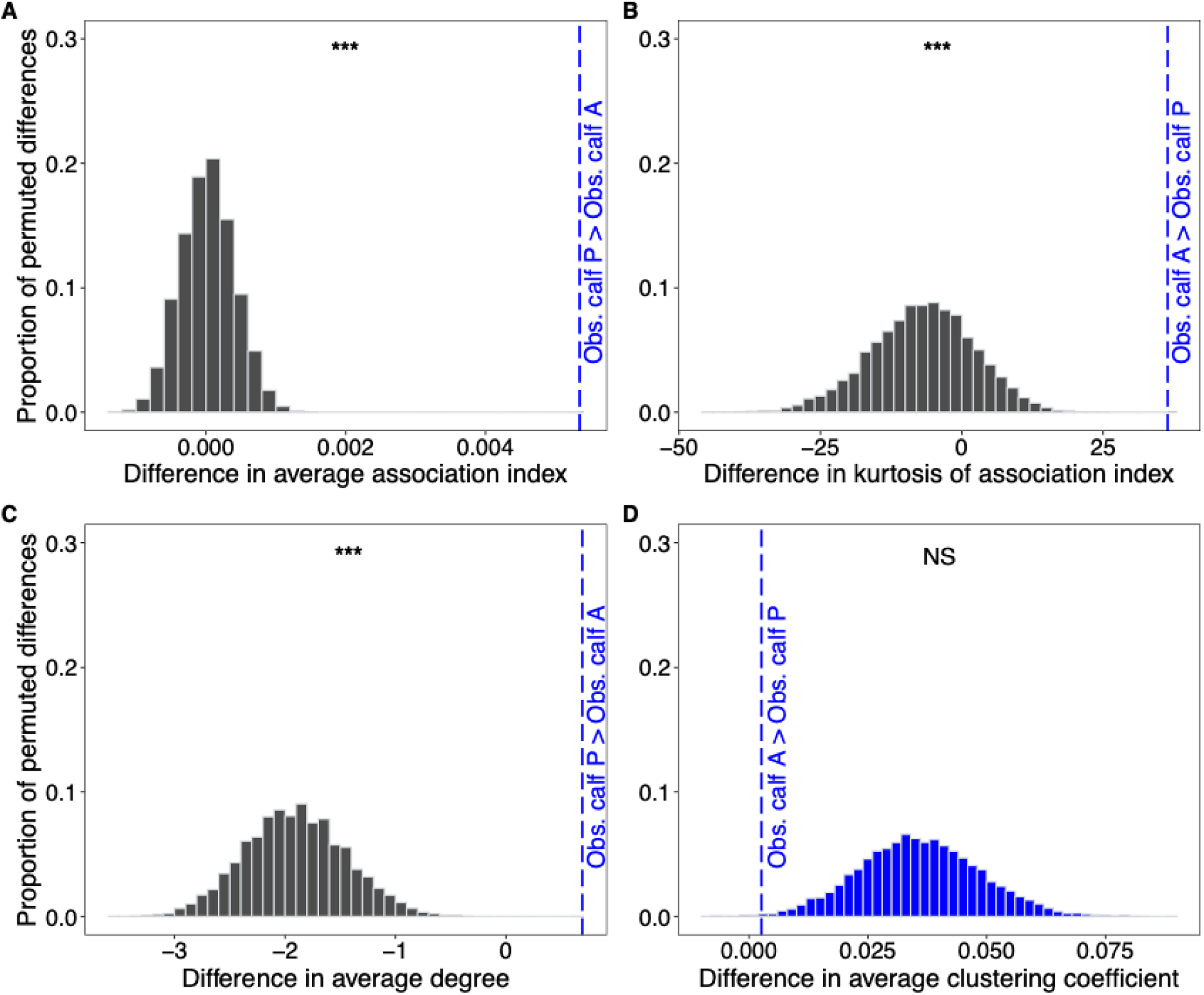
Differences in observed and permuted association index and network metrics of adult females. Distributions of the proportions of permuted datasets for difference in (A) average AI, (B) kurtosis of AI, (C) average degree, and (D) average clustering coefficient calculated from 10,000 permuted ‘calf-absent’ and ‘calf-present’ datasets, and the observed difference for each variable are shown. Differences from the randomised datasets that are at least as extreme as the observed difference are marked in blue. Significance codes: <0.001 ‘***’ 0.001 ‘**’ 0.01 ‘*’ 0.05 ‘.’ >0.05 ‘NS’.

Using sampled randomisation tests, we also found that the average degree was significantly higher in calf presence (12.5, SD=9.91) than absence (11.9, SD=9.89) (supplementary material, S6, table S1; Figure 3C). Average clustering coefficient was not significantly different between calf presence and absence (Figure 3D).

## 4. Discussion

In the first study of its kind, we examined whether the presence of calves affect female sociality in the Asian elephant, a species exhibiting a high degree of fission-fusion dynamics. We used an extensive dataset of about 5900 elephant group sightings collected over 9 years on hundreds of individually identified females of the Kabini elephant population [32,49,50,54]. Our results show that the number of females increased in the presence of calves in a group. Specifically, the number of younger age-class females without a dependent offspring of their own, who often provide allomaternal care in Asian elephants [24,26,47], increased in groups in calf presence. The pattern of results remained consistent when we investigated the number of females experienced by individual adult females in calf-present versus calf-absent groups. The presence of calves changed not just the social organisation but also the social structure of females. Our results show that adult females had a greater number of other adult females as associates, with whom they associated more strongly, in calf presence than absence. We discuss these findings below.

### (a) Calf protection and allomaternal care as reasons for an increase in female numbers in calf presence at both the group and individual level

Previously, McKay [40] had reported on Asian elephants in Sri Lanka that groups with calves had females apart from the mother, and that large groups (15 or more individuals), which were seen only occasionally, always had calves and juveniles. Being an early study, no further analysis was carried out to investigate how and why group size differences might have arisen. Larger group sizes in the presence of dependent offspring have also been found in a few other species that show fission-fusion dynamics (Bottlenose dolphins (*Tursiops truncatus*): [27,28,67]; Sumatran orangutans: [16], Ruffed lemurs (*Varecia variegata*): [12]; Warthogs (*Phacochoerus africanus*): [29]; Transient killer whales (*Orcinus orca*): [8]; Sperm whales (*Physeter macrocephalus*): [68]). However, with the exception of the results on a small subset of adult female orangutans [16], all other studies either examined group size as the number of individuals of all age-sex classes or had season confounding the presence of dependent offspring. In the Kabini elephant population, previous studies found no consistent group size variation within clans across seasons [14,51]. Therefore, our study provides the clearest positive effect of dependent offspring on female group size in a fission-fusion species, also see [51].

The differences in the magnitudes of increase in the numbers of females of the different age-classes offer insights into the potential reasons why calves have an effect on female social organisation. In the study population, and elsewhere in the wild and in captivity, Asian elephant dependent offspring have extremely high mortality rates [45,50,69,70] and face predation risk from tigers [44]. Therefore, the need for collective group defence of calves [46] is a likely explanation for the number of females being higher in calf-present than calf-absent groups. Despite clans having a greater number of adult females than young adult or subadult females (supplementary material, S1, table S1), we found that the magnitudes of increase of the numbers (and experienced numbers) of young adult and subadult females were greater than that of all adult females. This was also particularly true when we investigated only the numbers of adult females without their own dependent offspring. As females of these age-classes often provide allomaternal care in Asian as well as African savannah elephants (*Loxodonta Africana*) [26,71], our results strongly suggest that the need for allomaternal care is likely an important reason for the observed effect of calves on female social organisation. A caveat here is that although changes in the numbers of adult females in groups arise from fission-fusion dynamics, it is possible for the greater number of subadult females to partly (although not entirely) result from a demographic effect. Asian elephants have a long interbirth interval, which varies between 2.5-8 years in the wild [40, 46, 50, 72], but is often a minimum of 4 years [46]. Therefore, a calf and, even more likely, a young juvenile (≥1 and <2 years of age), will often be a subadult by the time its mother has had her next calf, possibly resulting in female subadults and calves being in the same group, if the female subadult continues to associate with her mother. As male Asian elephants do not provide parental care, the number of males in a group is unlikely to be strongly associated with the calf status predictor in our model (m_M), potentially explaining why the full model did not outperform the null model.

It is possible for variation in other factors that we have not examined, such as habitat type (forest vs grassland), food resource distribution (across zones within the grassland) [37], and predator distribution to affect group sizes. However, the majority of the sightings used for the current analyses was collected in the Kabini grassland area, which reduces the variation introduced by habitat type or predator distribution across sightings. Furthermore, calf-present or calf-absent sightings were not consistently limited to particular zones in the Kabini grassland. Due to denser vegetation and higher tiger density in forest than grassland, it is possible for the effect of calves on female group size to differ based on habitat type, such as that observed in giraffe [11]. However, because allomaternal care is crucial and frequent in Asian elephants across habitats [24,26,46,47], only the magnitude – and not the direction – of the effect of calves on female group composition and associations is likely to change based on habitat type.

### (b) Allomaternal care likely drives stronger associations among females in calf presence

Associations amongst females may be particularly useful for calf development and survival, through facilitating allomaternal care [26,40,45–47] as seen in a group of sperm whales [25]. Our results support this possibility. First, larger female group sizes in calf presence directly allowed for adult females to associate with a greater number of other adult females in their clans, see [31], leading to higher degree in calf presence than absence. Although the need for calf protection alone could have increased as well as strengthened associations, our results suggest that the need for allomaternal care is also partly responsible for driving changes in female association patterns.

The kurtosis of AI was greater than 3 (compared to that of a normal distribution) in both the presence and absence of calves, indicating that extreme values were more likely than in a normal distribution. The higher kurtosis of AI in calf absence than presence suggests than higher AI values are an outlier (in relation to the distribution) in calf absence, whereas they are not substantial outliers in calf presence. Indeed, there were a larger number of high AI values in the presence than absence of calves. Specifically, there was a higher percentage of adult females with association index values of over 0.5, pointing to strengthened associations in calf presence. We previously found that the same female often allomothered a calf, and many of the allomothers almost always associated with the calves they allomothered (and the calves’ mothers) throughout at least the first year of the calf’s life [26]. Therefore, the strengthened associations are likely at least partly driven by the associations between mothers and allomothers in calf presence. We also found a larger number of associates (degree) of adult females in the social network in calf presence than absence, indicating more connections in calf presence. Clustering coefficient was not significantly different between calf presence and absence, but this is likely a result of a large number of connections within clans being realised even in calf absence. Degree and association strength were higher during lactation than mating period in female degus [30]. Currently, we do not have comparable results on any other species for these metrics. Thus, our study shows the positive effect of calves on multiple aspects of female social structure.

### (c) Context-dependent effect of dependent offspring on female sociality in fission-fusion species

Although our study reveals a strong, positive effect of the presence of calves on female sociality, in other fission-fusion species, depending on the ecological and social pressures that have shaped their social systems, this effect may be attenuated or may even be reversed. Due to the absence of female-driven infanticide in elephants [41], the emergence of cooperation among females in the context of care has become feasible. In contrast, the presence of female-driven infanticide risk reduces rather than increases female sociality in the presence of dependent offspring, such as in spotted hyenas [7]. Furthermore, prevailing ecological conditions may influence whether increased female sociality can be expressed in dependent offspring presence. For example, depending on habitat type and predator density, females with dependent offspring may either form large or small groups. In open habitats with low predation risk, females may rely on increased vigilance and better predator detection and avoidance, whereas in dense habitats with high predation risk females may adopt alternative strategies to mitigate the risk, such as giraffes hiding their calves when in dense habitats with high lion density [11]. Further studies on other populations and species with varied socioecological pressures and prevailing conditions are needed to fully understand the factors deciding the magnitude and the direction of the effect of dependent offspring on female sociality in fission-fusion species.

### (d) Future work

Our study opens up several avenues for future research. First, if female sociality increased in order to enhance offspring protection and/or allomaternal care, we should expect to find a decrease in the effect of offspring presence on female sociality with offspring age as their survival probability improves, such as that seen in dolphins [27]. Additionally, increase in group size and associations should come with benefits for the dependent offspring. Second, increased female association strength in the presence of dependent offspring may also arise from mothers preferentially associating with one another due to shared nutritional needs, as seen in Bechstein’s bat (*Myotis bechsteinii*) [73] and mountain goats (*Oreamnos americanus*) [74]. This could be examined in the future. Third, a non-mutually exclusive explanation for preferential associations among mothers with similarly-aged offspring could be the need to enhance opportunities for peer socialisation and play as recently demonstrated in Bornean orangutans (*Pongo pygmaeus wurmbii*) [75]. Finally, although an increase in female sociality may carry benefits, there may also be costs, see [31,51], such as increased feeding competition.

## 5. Conclusion

Our work provides the first direct and concrete evidence that calves increase female sociality in Asian elephants exhibiting a high degree of fission-fusion dynamics. As argued by a few others before us [47,71], given the significant role played by dependent offspring in altering female Asian elephant social organisation and structure, we posit that dependent offspring may have been a factor important in the evolution of female social relationships in this species. When considered together with direct and indirect evidence from other fission-fusion species [8,12,16,25,27–30,67,68,75], our study makes it clear that the need to protect dependent offspring and to provide allomaternal care can bring together females and strengthen female social bonds, conditional upon prevailing conditions and the socioecological pressures that have shaped the social system of the species.

## Ethics

Field data collection involved observations of wild elephants at a minimum distance of ∼40 m so as to minimise any disturbance to their natural behaviour. The study was purely observational and non-invasive in nature and did not involve any animal handling or manipulation. Fieldwork was carried out under permit numbers D/WL/CR-121/2008–2009 dated 23/01/09, 18/05/10, and 08/05/12, and PCCF(WL)/E2/CR-121/2014-15 dated 09/05/14 and 28/05/2019, issued by the office of the Principal Chief Conservator of Forests (Wildlife) and Chief Wildlife Warden, Karnataka.

## Declaration of AI use

We have not used AI-assisted technologies in creating this article.

## Acknowledgements

We thank the offices of the Principal Chief Conservator of Forests - Wildlife (PCCF-WL), Karnataka Forest Department, and of the Conservators, Deputy Conservators of Forests, and Field Directors of Bandipur and Nagarahole National Parks and Tiger Reserves for field permits. We also thank various officials, from the PCCF(WL) and Additional PCCF(WL), to the Conservators of Forests, Deputy Conservators of Forests, Assistant Conservators of Forests, and Range Forest Officers, to the staff of Nagarahole and Bandipur National Parks and Tiger Reserves for their support over the years. We thank our field trackers Gunda, Rajesh, Krishna, Shankar, and Ranga, drivers Binu, Althaf, and Pramod, and others for field assistance. We thank Deepika Prasad and Arjun Ghosh for helping with initial data collection. This work is part of the Ph.D. thesis of TR.

