## Supplemental Information for "Calves function as social magnets in a species with high fission-fusion dynamics, the Asian elephant"

Supplementary Material S1.

*Clan demography*

**Supplementary Material S1, Table S1**. Variation in clan demography across the clans used for the group size, experienced group size, and association analyses. The clans used for the three different analyses varied (*: used for group size analysis; †: used for experienced group size analysis; all the 18 clans were used for association analysis). For each clan, the number of adult females (≥10 years; white rows), young adult females (≥10 and <15 years; green rows), and subadult females (≥5 and <10 years; grey rows) in each year are provided, along with the mean (SD) over the entire study period. Note that the clans were not sighted across all the sampling years (2009-2018).

| Clan/Year | 2009 | 2010 | 2011 | 2012 | 2013 | 2014 | 2015 | 2016 | 2017 | 2018 | Mean (SD) |
| --- | --- | --- | --- | --- | --- | --- | --- | --- | --- | --- | --- |
|  | 11 | 11 | 11 | 10 | 9 | 9 | 9 | 10 | 8 | 8 | 9.6 (1.17) |
| Alexandra*^†^ | 3 | 2 | 2 | 1 | 1 | 0 | 0 | 1 | 1 | 1 | 1.2 (0.92) |
|  | 0 | 0 | 1 | 2 | 2 | 2 | 1 | 0 | 0 | 0 | 0.8 (0.92) |
|  | 11 | 11 | 11 | 9 | 8 | 10 | 10 | 11 | 12 | 11 | 10.4 (1.17) |
| Anabelle* | 4 | 3 | 3 | 1 | 0 | 2 | 2 | 4 | 5 | 5 | 2.9 (1.66) |
|  | 2 | 2 | 4 | 5 | 5 | 3 | 3 | 3 | 2 | 2 | 3.1 (1.20) |
|  | 7 | 7 | 6 | 6 | 6 | 6 | 6 | 5 | 5 | 5 | 5.9 (0.74) |
| Cleopatra | 1 | 1 | 1 | 0 | 0 | 0 | 0 | 0 | 1 | 2 | 0.6 (0.70) |
|  | 0 | 0 | 0 | 1 | 2 | 2 | 2 | 2 | 1 | 0 | 1.0 (0.94) |
|  | 5 | 5 | 7 | 7 | 7 | 9 | 10 | 10 | 10 | 10 | 8.0 (2.05) |
| Fiola | 1 | 1 | 3 | 3 | 3 | 4 | 5 | 3 | 3 | 3 | 2.9 (1.20) |
|  | 4 | 5 | 3 | 3 | 3 | 1 | 0 | 1 | 1 | 1 | 2.2 (1.62) |
|  | 4 | 5 | 5 | 5 | 5 | 5 | 6 | 6 | 7 | 7 | 5.5 (0.97) |
| Jyena | 1 | 1 | 1 | 1 | 1 | 1 | 1 | 1 | 2 | 2 | 1.2 (0.42) |
|  | 1 | 1 | 1 | 2 | 2 | 2 | 1 | 1 | 0 | 0 | 1.1 (0.74) |
|  | 7 | 8 | 7 | 7 | 7 | 7 | 7 | 7 | 8 | 9 | 7.4 (0.70) |
| Kasturi*^†^ | 0 | 1 | 1 | 1 | 1 | 1 | 0 | 0 | 1 | 2 | 0.8 (0.63) |
|  | 1 | 0 | 0 | 1 | 2 | 2 | 2 | 2 | 1 | 0 | 1.1 (0.88) |
|  | 16 | 16 | 16 | 14 | 13 | 15 | 16 | 15 | 14 | 14 | 14.9 (1.10) |
| Katrina* | 4 | 3 | 3 | 0 | 0 | 2 | 3 | 3 | 3 | 3 | 2.4 (1.35) |
|  | 2 | 3 | 3 | 3 | 3 | 1 | 0 | 0 | 0 | 0 | 1.5 (1.43) |
|  | 4 | 4 | 4 | 4 | 3 | 3 | 3 | 3 | 4 | 5 | 3.7 (0.67) |
| Lalanti | 0 | 0 | 0 | 0 | 0 | 0 | 0 | 0 | 1 | 2 | 0.3 (0.67) |
|  | 0 | 0 | 0 | 1 | 2 | 2 | 2 | 2 | 1 | 0 | 1.0 (0.94) |
|  | 16 | 16 | 16 | 15 | 15 | 14 | 15 | 15 | 12 | 12 | 14.6 (1.51) |
| Lisa*^†^ | 3 | 2 | 2 | 1 | 2 | 2 | 4 | 3 | 3 | 2 | 2.4 (0.84) |
|  | 2 | 4 | 3 | 3 | 2 | 3 | 0 | 0 | 0 | 0 | 1.7 (1.57) |
|  | 9 | 9 | 9 | 10 | 10 | 10 | 10 | 10 | 8 | 10 | 9.5 (0.71) |
| Menaka* | 2 | 1 | 1 | 2 | 2 | 1 | 1 | 1 | 0 | 2 | 1.3 (0.67) |
|  | 1 | 1 | 1 | 0 | 2 | 2 | 2 | 2 | 2 | 0 | 1.3 (0.82) |
|  | 6 | 6 | 6 | 7 | 7 | 9 | 9 | 9 | 9 | 9 | 7.7 (1.42) |
| Mridula* | 1 | 0 | 0 | 1 | 1 | 3 | 3 | 4 | 3 | 3 | 1.9 (1.45) |
|  | 3 | 3 | 4 | 3 | 3 | 1 | 1 | 0 | 0 | 1 | 1.9 (1.45) |
|  | 14 | 15 | 14 | 14 | 14 | 14 | 16 | 16 | 14 | 14 | 14.5 (0.85) |
| Nakshatra*^†^ | 3 | 4 | 4 | 3 | 3 | 2 | 2 | 2 | 3 | 4 | 3.0 (0.82) |
|  | 2 | 2 | 2 | 3 | 4 | 4 | 2 | 2 | 1 | 0 | 2.2 (1.23) |
|  | 23 | 23 | 23 | 24 | 25 | 24 | 26 | 25 | 26 | 28 | 24.7 (1.64) |
| Olympia* | 4 | 3 | 5 | 3 | 5 | 4 | 7 | 5 | 5 | 6 | 4.7 (1.25) |
|  | 5 | 8 | 6 | 6 | 6 | 7 | 4 | 4 | 3 | 2 | 5.1 (1.85) |
|  | 19 | 17 | 17 | 16 | 15 | 16 | 17 | 19 | 20 | 20 | 17.6 (1.78) |
| Osanna* | 4 | 3 | 3 | 2 | 2 | 2 | 3 | 4 | 6 | 6 | 3.5 (1.51) |
|  | 2 | 3 | 4 | 6 | 6 | 5 | 6 | 5 | 2 | 2 | 4.1 (1.73) |
|  | 20 | 21 | 21 | 19 | 19 | 18 | 20 | 23 | 23 | 24 | 20.8 (1.99) |
| Patricia*^†^ | 5 | 5 | 5 | 3 | 4 | 2 | 3 | 6 | 8 | 8 | 4.9 (2.02) |
|  | 2 | 4 | 7 | 10 | 10 | 9 | 7 | 4 | 1 | 0 | 5.4 (3.72) |
|  | 4 | 4 | 4 | 5 | 5 | 6 | 6 | 6 | 6 | 4 | 5.0 (0.94) |
| Tilottama* | 0 | 0 | 0 | 1 | 1 | 2 | 2 | 2 | 1 | 1 | 1.0 (0.82) |
|  | 2 | 2 | 2 | 1 | 1 | 0 | 0 | 0 | 0 | 0 | 0.8 (0.92) |
|  | 13 | 14 | 13 | 13 | 14 | 14 | 14 | 15 | 15 | 18 | 14.3 (1.49) |
| Unnati* | 1 | 2 | 2 | 1 | 2 | 2 | 1 | 2 | 2 | 4 | 1.9 (0.88) |
|  | 2 | 1 | 2 | 2 | 4 | 5 | 6 | 6 | 6 | 3 | 3.7 (1.95) |
|  | 30 | 29 | 30 | 27 | 29 | 32 | 34 | 35 | 33 | 35 | 31.4 (2.80) |
| Victoria*^†^ | 8 | 7 | 7 | 7 | 9 | 8 | 10 | 11 | 12 | 12 | 9.1 (2.02) |
|  | 8 | 11 | 12 | 15 | 17 | 16 | 14 | 14 | 7 | 7 | 12.1 (3.73) |

As clan demography clearly differed across the clans used for analyses, we controlled for it in our models. For more details about the analysis, refer to the data analysis section of the Methods.

Supplementary Material S2.

*Sample size for the group size and the experienced group size analyses*

**Supplementary Material S2, Table S1**. Clan sampling for the analysis on the number of females of different age-classes in a group. Number of sightings in calf absence (white rows) and presence (green rows) are shown, along with the total number of sightings for each clan in calf absence and presence and for each year. “_” means that a particular clan was not observed at least five times in either calf absence or presence (or both) in a particular year.

| Clan/Year | 2009 | 2010 | 2011 | 2012 | 2013 | 2014 | 2015 | 2016 | 2017 | 2018 | **Total** |
| --- | --- | --- | --- | --- | --- | --- | --- | --- | --- | --- | --- |
| Alexandra | _ | _ | _ | _ | 12 | _ | 11 | 13 | _ | 8 | 44 |
|  | _ | _ | _ | _ | 6 | _ | 15 | 19 | _ | 10 | 50 |
| Anabelle | _ | _ | 16 | _ | _ | _ | _ | _ | _ | _ | 16 |
|  | _ | _ | 15 | _ | _ | _ | _ | _ | _ | _ | 15 |
| Kasturi | 30 | _ | 56 | 47 | _ | _ | _ | 34 | _ | 35 | 202 |
|  | 7 | _ | 7 | 57 | _ | _ | _ | 46 | _ | 55 | 172 |
| Katrina | _ | _ | _ | _ | 7 | _ | 11 | _ | 6 | _ | 24 |
|  | _ | _ | _ | _ | 8 | _ | 6 | _ | 13 | _ | 27 |
| Lisa | 15 | _ | 44 | 107 | 91 | 50 | 35 | 80 | 44 | _ | 466 |
|  | 7 | _ | 12 | 24 | 11 | 18 | 7 | 54 | 21 | _ | 154 |
| Menaka | _ | _ | _ | _ | 19 | _ | _ | 14 | _ | _ | 33 |
|  | _ | _ | _ | _ | 5 | _ | _ | 25 | _ | _ | 30 |
| Mridula | _ | _ | _ | _ | 24 | _ | 12 | _ | _ | _ | 36 |
|  | _ | _ | _ | _ | 5 | _ | 5 | _ | _ | _ | 10 |
| Nakshatra | 10 | _ | 19 | _ | 30 | 22 | _ | 61 | _ | 35 | 177 |
|  | 21 | _ | 7 | _ | 13 | 23 | _ | 39 | _ | 16 | 119 |
| Olympia | _ | _ | 16 | 17 | 36 | _ | _ | _ | _ | _ | 69 |
|  | _ | _ | 14 | 17 | 11 | _ | _ | _ | _ | _ | 42 |
| Osanna | _ | _ | 6 | _ | _ | _ | 11 | _ | 18 | _ | 35 |
|  | _ | _ | 12 | _ | _ | _ | 6 | _ | 6 | _ | 24 |
| Patricia | _ | 25 | 106 | 144 | _ | _ | 167 | 228 | 52 | 71 | 793 |
|  | _ | 8 | 6 | 38 | _ | _ | 47 | 8 | 51 | 39 | 197 |
| Tilottama | _ | _ | 5 | _ | 5 | _ | _ | 5 | _ | _ | 15 |
|  | _ | _ | 11 | _ | 11 | _ | _ | 38 | _ | _ | 60 |
| Unnati | _ | _ | 6 | _ | _ | _ | _ | _ | _ | _ | 6 |
|  | _ | _ | 7 | _ | _ | _ | _ | _ | _ | _ | 7 |
| Victoria | 76 | _ | 49 | 173 | 84 | 119 | _ | 61 | _ | 77 | 639 |
|  | 19 | _ | 80 | 28 | 64 | 106 | _ | 169 | _ | 40 | 506 |
| **Total** | 185 | 33 | 494 | 652 | 442 | 338 | 333 | 894 | 211 | 386 | **3968** |

**Supplementary Material S2, Table S2**. Adult female sampling for the experienced number of females analysis. The number of sightings for each female in both calf absence (white rows) and presence (green rows) within each year for each clan are shown. Only females who were sighted at least five times were chosen for the analyses. “_” means that a particular adult female was not observed for at least five times in either calf absence or presence (or both) in a particular year. Note that we could not add data from 2010 because in none of the sampled clans we had at least five sightings of an adult female in both calf absence and presence.

| **Clan ID: Alexandra** | | | | | | | | | | | |
| --- | --- | --- | --- | --- | --- | --- | --- | --- | --- | --- | --- |
| Female ID/Year | 2009 | 2010 | 2011 | 2012 | 2013 | 2014 | 2015 | 2016 | 2017 | 2018 | **Total** |
| Aleesha | _ | _ | _ | _ | _ | _ | _ | 6 | _ | _ | 6 |
|  | _ | _ | _ | _ | _ | _ | _ | 9 | _ | _ | 9 |
| Alena | _ | _ | _ | _ | _ | _ | 6 | _ | _ | _ | 6 |
|  | _ | _ | _ | _ | _ | _ | 8 | _ | _ | _ | 8 |
| Zarin | _ | _ | _ | _ | _ | _ | 6 | _ | _ | _ | 6 |
|  | _ | _ | _ | _ | _ | _ | 11 | _ | _ | _ | 11 |
| Zohra | _ | _ | _ | _ | _ | _ | 8 | _ | _ | _ | 8 |
|  | _ | _ | _ | _ | _ | _ | 7 | _ | _ | _ | 7 |
| Zoya | _ | _ | _ | _ | _ | _ | _ | 9 | _ | _ | 9 |
|  | _ | _ | _ | _ | _ | _ | _ | 7 | _ | _ | 7 |

| **Clan ID: Kasturi** | | | | | | | | | | | |
| --- | --- | --- | --- | --- | --- | --- | --- | --- | --- | --- | --- |
| Female ID/Year | 2009 | 2010 | 2011 | 2012 | 2013 | 2014 | 2015 | 2016 | 2017 | 2018 | **Total** |
| Kasturi | _ | _ | _ | 8 | _ | _ | _ | 29 | _ | 12 | 49 |
|  | _ | _ | _ | 46 | _ | _ | _ | 15 | _ | 28 | 89 |
| Kausalya | _ | _ | _ | 15 | _ | _ | _ | _ | _ | 19 | 34 |
|  | _ | _ | _ | 40 | _ | _ | _ | _ | _ | 27 | 67 |
| Kausavi | _ | _ | _ | _ | _ | _ | _ | _ | _ | 17 | 17 |
|  | _ | _ | _ | _ | _ | _ | _ | _ | _ | 26 | 26 |
| Keerthana | _ | _ | _ | 6 | _ | _ | _ | _ | _ | _ | 6 |
|  | _ | _ | _ | 31 | _ | _ | _ | _ | _ | _ | 31 |
| Ketki | _ | _ | _ | 22 | _ | _ | _ | _ | _ | 10 | 32 |
|  | _ | _ | _ | 25 | _ | _ | _ | _ | _ | 33 | 58 |
| Kokila | _ | _ | _ | 18 | _ | _ | _ | _ | _ | 12 | 30 |
|  | _ | _ | _ | 26 | _ | _ | _ | _ | _ | 27 | 53 |
| Koyna | _ | _ | 10 | 6 | _ | _ | _ | _ | _ | 5 | 21 |
|  | _ | _ | 5 | 32 | _ | _ | _ | _ | _ | 37 | 74 |

| **Clan ID: Lisa** | | | | | | | | | | | |
| --- | --- | --- | --- | --- | --- | --- | --- | --- | --- | --- | --- |
| Female ID/Year | 2009 | 2010 | 2011 | 2012 | 2013 | 2014 | 2015 | 2016 | 2017 | 2018 | **Total** |
| Jacintha | _ | _ | _ | 9 | _ | _ | _ | 10 | _ | _ | 19 |
|  | _ | _ | _ | 10 | _ | _ | _ | 10 | _ | _ | 20 |
| Joan | _ | _ | _ | 12 | _ | _ | _ | _ | _ | _ | 12 |
|  | _ | _ | _ | 11 | _ | _ | _ | _ | _ | _ | 11 |
| Joycee | _ | _ | _ | 11 | _ | _ | _ | _ | _ | _ | 11 |
|  | _ | _ | _ | 10 | _ | _ | _ | _ | _ | _ | 10 |
| Komal | _ | _ | _ | _ | _ | _ | _ | 9 | _ | _ | 9 |
|  | _ | _ | _ | _ | _ | _ | _ | 33 | _ | _ | 33 |
| Laika | _ | _ | _ | _ | _ | 5 | _ | 9 | _ | _ | 14 |
|  | _ | _ | _ | _ | _ | 11 | _ | 18 | _ | _ | 29 |
| Lana | _ | _ | _ | _ | _ | _ | _ | 11 | 7 | _ | 18 |
|  | _ | _ | _ | _ | _ | _ | _ | 28 | 7 | _ | 35 |
| Laura | _ | _ | _ | 15 | _ | 7 | _ | 11 | 6 | _ | 39 |
|  | _ | _ | _ | 5 | _ | 13 | _ | 17 | 11 | _ | 46 |
| Lauren | _ | _ | _ | _ | _ | _ | _ | 9 | _ | _ | 9 |
|  | _ | _ | _ | _ | _ | _ | _ | 25 | _ | _ | 25 |
| Leena | _ | _ | _ | 11 | _ | 12 | _ | 19 | _ | _ | 42 |
|  | _ | _ | _ | 11 | _ | 12 | _ | 23 | _ | _ | 46 |
| Liezl | _ | _ | _ | 20 | _ | 19 | 5 | 20 | _ | _ | 64 |
|  | _ | _ | _ | 7 | _ | 6 | 7 | 14 | _ | _ | 34 |
| Lily | 6 | _ | _ | 21 | _ | 18 | _ | 36 | 23 | _ | 104 |
|  | 5 | _ | _ | 5 | _ | 6 | _ | 15 | 7 | _ | 38 |
| Linda | _ | _ | _ | 13 | _ | _ | _ | _ | _ | _ | 13 |
|  | _ | _ | _ | 11 | _ | _ | _ | _ | _ | _ | 11 |
| Linette | _ | _ | _ | 25 | _ | _ | _ | _ | _ | _ | 25 |
|  | _ | _ | _ | 10 | _ | _ | _ | _ | _ | _ | 10 |
| Lydia | _ | _ | _ | 18 | 13 | 8 | _ | 16 | 7 | _ | 62 |
|  | _ | _ | _ | 8 | 7 | 12 | _ | 33 | 14 | _ | 74 |
| Lynn | _ | _ | _ | 20 | _ | 5 | 10 | 9 | _ | _ | 44 |
|  | _ | _ | _ | 5 | _ | 17 | 7 | 41 | _ | _ | 70 |

| **Clan ID: Patricia** | | | | | | | | | | | |
| --- | --- | --- | --- | --- | --- | --- | --- | --- | --- | --- | --- |
| Female ID/Year | 2009 | 2010 | 2011 | 2012 | 2013 | 2014 | 2015 | 2016 | 2017 | 2018 | **Total** |
| Candice | _ | _ | _ | _ | _ | _ | 13 | _ | _ | _ | 13 |
|  | _ | _ | _ | _ | _ | _ | 9 | _ | _ | _ | 9 |
| Casey | _ | _ | _ | _ | _ | _ | 5 | _ | _ | _ | 5 |
|  | _ | _ | _ | _ | _ | _ | 28 | _ | _ | _ | 28 |
| Cassandra | _ | _ | _ | 8 | _ | _ | 6 | 31 | _ | _ | 45 |
|  | _ | _ | _ | 5 | _ | _ | 29 | 5 | _ | _ | 39 |
| Dana | _ | _ | _ | 18 | _ | _ | _ | _ | _ | _ | 18 |
|  | _ | _ | _ | 28 | _ | _ | _ | _ | _ | _ | 28 |
| Dominique | _ | _ | _ | _ | _ | _ | _ | _ | _ | 10 | 10 |
|  | _ | _ | _ | _ | _ | _ | _ | _ | _ | 8 | 8 |
| Dorothy | _ | _ | _ | _ | _ | _ | _ | _ | _ | 8 | 8 |
|  | _ | _ | _ | _ | _ | _ | _ | _ | _ | 8 | 8 |
| Gemini | _ | _ | _ | _ | _ | _ | _ | _ | _ | 8 | 8 |
|  | _ | _ | _ | _ | _ | _ | _ | _ | _ | 8 | 8 |
| Genette | _ | _ | _ | _ | _ | _ | _ | _ | 14 | _ | 14 |
|  | _ | _ | _ | _ | _ | _ | _ | _ | 8 | _ | 8 |
| Genevieve | _ | _ | _ | _ | _ | _ | _ | _ | _ | 6 | 6 |
|  | _ | _ | _ | _ | _ | _ | _ | _ | _ | 23 | 23 |
| Georgetta | _ | _ | _ | _ | _ | _ | _ | _ | _ | 8 | 8 |
|  | _ | _ | _ | _ | _ | _ | _ | _ | _ | 8 | 8 |
| Georgina | _ | _ | _ | _ | _ | _ | _ | _ | _ | 8 | 8 |
|  | _ | _ | _ | _ | _ | _ | _ | _ | _ | 8 | 8 |
| Glenda | _ | _ | _ | _ | _ | _ | 13 | _ | _ | _ | 13 |
|  | _ | _ | _ | _ | _ | _ | 5 | _ | _ | _ | 5 |
| Pamela | _ | _ | _ | _ | _ | _ | 14 | 35 | _ | _ | 49 |
|  | _ | _ | _ | _ | _ | _ | 23 | 5 | _ | _ | 28 |
| Pandora | _ | _ | _ | _ | _ | _ | _ | 33 | _ | _ | 33 |
|  | _ | _ | _ | _ | _ | _ | _ | 5 | _ | _ | 5 |
| Patricia | _ | _ | _ | _ | _ | _ | 20 | 24 | _ | _ | 44 |
|  | _ | _ | _ | _ | _ | _ | 21 | 5 | _ | _ | 26 |

| **Clan ID: Victoria** | | | | | | | | | | | |
| --- | --- | --- | --- | --- | --- | --- | --- | --- | --- | --- | --- |
| Female ID/Year | 2009 | 2010 | 2011 | 2012 | 2013 | 2014 | 2015 | 2016 | 2017 | 2018 | **Total** |
| Fanaa | _ | _ | _ | _ | _ | _ | _ | _ | _ | 6 | 6 |
|  | _ | _ | _ | _ | _ | _ | _ | _ | _ | 5 | 5 |
| Fawzia | _ | _ | 5 | _ | _ | _ | _ | _ | _ | _ | 5 |
|  | _ | _ | 5 | _ | _ | _ | _ | _ | _ | _ | 5 |
| Floppy_ears | _ | _ | _ | _ | 5 | 13 | _ | _ | _ | _ | 18 |
|  | _ | _ | _ | _ | 13 | 17 | _ | _ | _ | _ | 30 |
| Impedimenta | _ | _ | 15 | _ | _ | 10 | _ | 8 | _ | _ | 33 |
|  | _ | _ | 6 | _ | _ | 28 | _ | 46 | _ | _ | 80 |
| Ina | 21 | _ | _ | _ | _ | 11 | _ | 6 | _ | _ | 38 |
|  | 5 | _ | _ | _ | _ | 22 | _ | 13 | _ | _ | 40 |
| Ipomoea | 7 | _ | 6 | _ | _ | 17 | _ | 14 | _ | _ | 44 |
|  | 14 | _ | 7 | _ | _ | 8 | _ | 39 | _ | _ | 68 |
| Itamara | 28 | _ | 7 | _ | _ | 24 | _ | 7 | _ | 5 | 71 |
|  | 6 | _ | 11 | _ | _ | 22 | _ | 25 | _ | 11 | 75 |
| Saina | 23 | _ | _ | _ | _ | _ | _ | _ | _ | _ | 23 |
|  | 9 | _ | _ | _ | _ | _ | _ | _ | _ | _ | 9 |
| Sajida | _ | _ | _ | _ | 7 | 23 | _ | 10 | _ | 9 | 49 |
|  | _ | _ | _ | _ | 10 | 27 | _ | 12 | _ | 10 | 59 |
| Salma | _ | _ | _ | _ | _ | 15 | _ | _ | _ | 10 | 25 |
|  | _ | _ | _ | _ | _ | 19 | _ | _ | _ | 6 | 25 |
| Salvia | _ | _ | _ | _ | 9 | 16 | _ | _ | _ | _ | 25 |
|  | _ | _ | _ | _ | 14 | 19 | _ | _ | _ | _ | 33 |
| Sameena | _ | _ | 9 | _ | _ | _ | _ | _ | _ | _ | 9 |
|  | _ | _ | 9 | _ | _ | _ | _ | _ | _ | _ | 9 |
| Sarah | 21 | _ | 5 | _ | _ | _ | _ | 5 | _ | _ | 31 |
|  | 6 | _ | 9 | _ | _ | _ | _ | 13 | _ | _ | 28 |
| Sarayu | _ | _ | _ | _ | _ | 12 | _ | _ | _ | _ | 12 |
|  | _ | _ | _ | _ | _ | 16 | _ | _ | _ | _ | 16 |
| Sarin | 23 | _ | 5 | _ | 7 | 8 | _ | _ | _ | _ | 43 |
|  | 6 | _ | 12 | _ | 6 | 24 | _ | _ | _ | _ | 48 |
| Serena | _ | _ | _ | _ | 5 | 18 | _ | _ | _ | 9 | 32 |
|  | _ | _ | _ | _ | 9 | 25 | _ | _ | _ | 9 | 43 |
| Sunaina | 6 | _ | 7 | _ | _ | 9 | _ | _ | _ | _ | 22 |
|  | 6 | _ | 11 | _ | _ | 11 | _ | _ | _ | _ | 28 |
| Suneeti | _ | _ | _ | _ | _ | 9 | _ | _ | _ | _ | 9 |
|  | _ | _ | _ | _ | _ | 10 | _ | _ | _ | _ | 10 |
| Sunetra | _ | _ | 5 | _ | _ | 11 | _ | 5 | _ | _ | 21 |
|  | _ | _ | 7 | _ | _ | 20 | _ | 26 | _ | _ | 53 |
| Supri | _ | _ | _ | _ | _ | 5 | _ | 6 | _ | 10 | 21 |
|  | _ | _ | _ | _ | _ | 18 | _ | 24 | _ | 5 | 47 |
| Surabhi | _ | _ | _ | _ | _ | 9 | _ | _ | _ | _ | 9 |
|  | _ | _ | _ | _ | _ | 7 | _ | _ | _ | _ | 7 |
| Sushma | _ | _ | _ | _ | _ | 8 | _ | _ | _ | _ | 8 |
|  | _ | _ | _ | _ | _ | 8 | _ | _ | _ | _ | 8 |
| Suvantika | _ | _ | _ | _ | _ | _ | _ | _ | _ | 19 | 19 |
|  | _ | _ | _ | _ | _ | _ | _ | _ | _ | 5 | 5 |
| Suvarna | _ | _ | _ | _ | _ | _ | _ | 6 | _ | _ | 6 |
|  | _ | _ | _ | _ | _ | _ | _ | 70 | _ | _ | 70 |
| Suvarti | 8 | _ | 10 | 9 | _ | _ | _ | 10 | _ | _ | 37 |
|  | 6 | _ | 10 | 13 | _ | _ | _ | 28 | _ | _ | 57 |
| Suveera | _ | _ | _ | _ | 10 | _ | _ | _ | _ | _ | 10 |
|  | _ | _ | _ | _ | 19 | _ | _ | _ | _ | _ | 19 |
| Suvrata | _ | _ | _ | _ | _ | _ | _ | _ | _ | 5 | 5 |
|  | _ | _ | _ | _ | _ | _ | _ | _ | _ | 17 | 17 |
| Ursula | _ | _ | 5 | _ | _ | 15 | _ | _ | _ | _ | 20 |
|  | _ | _ | 6 | _ | _ | 7 | _ | _ | _ | _ | 13 |
| Valerie | 10 | _ | _ | 31 | _ | 19 | _ | _ | _ | _ | 60 |
|  | 5 | _ | _ | 8 | _ | 28 | _ | _ | _ | _ | 41 |
| Vanessa | 15 | _ | _ | _ | 18 | 10 | _ | 5 | _ | _ | 48 |
|  | 6 | _ | _ | _ | 8 | 26 | _ | 70 | _ | _ | 110 |
| Vera | 12 | _ | _ | 13 | _ | _ | _ | _ | _ | _ | 25 |
|  | 6 | _ | _ | 5 | _ | _ | _ | _ | _ | _ | 11 |
| Vicky | _ | _ | _ | _ | _ | _ | _ | 10 | _ | _ | 10 |
|  | _ | _ | _ | _ | _ | _ | _ | 68 | _ | _ | 68 |
| Victoria | 13 | _ | _ | _ | _ | _ | _ | _ | _ | _ | 13 |
|  | 5 | _ | _ | _ | _ | _ | _ | _ | _ | _ | 5 |

| **All females across all clans** | | | | | | | | | | | |
| --- | --- | --- | --- | --- | --- | --- | --- | --- | --- | --- | --- |
| Year | 2009 | 2010 | 2011 | 2012 | 2013 | 2014 | 2015 | 2016 | 2017 | 2018 | **Total** |
| No. of sightings | 278 | _ | 211 | 681 | 189 | 820 | 261 | 1347 | 104 | 530 | **4421** |

**Supplementary Material S2, Table S3**. Adult female sampling for the association analysis. The number of sightings for each female in calf absence and presence are provided. Each female was sighted at least five times in both calf presence and absence over the entire sampling period. The range of the number of sightings of adult females in calf absence was 5-274 and that in calf presence was 5-169. The total number of adult females used for the analysis was 171. The number of sampled adult females per clan ranged from 2 to 35.

| SI No | Female ID | Clan ID | Calf absence | Calf presence | **Total no. of times sighted** |
| --- | --- | --- | --- | --- | --- |
| 1 | Aleesha | Alexandra | 25 | 27 | 52 |
| 2 | Alena | Alexandra | 17 | 36 | 53 |
| 3 | Alesti | Alexandra | 12 | 6 | 18 |
| 4 | Althea | Alexandra | 29 | 13 | 42 |
| 5 | Zarin | Alexandra | 20 | 40 | 60 |
| 6 | Zerad | Alexandra | 14 | 21 | 35 |
| 7 | Zhaleh | Alexandra | 12 | 35 | 47 |
| 8 | Zohra | Alexandra | 19 | 26 | 45 |
| 9 | Zoya | Alexandra | 28 | 29 | 57 |
| 10 | Anca | Anabelle | 39 | 14 | 53 |
| 11 | Andrea | Anabelle | 62 | 12 | 74 |
| 12 | Annette | Anabelle | 29 | 13 | 42 |
| 13 | Antonia | Anabelle | 57 | 12 | 69 |
| 14 | Wanda | Anabelle | 74 | 14 | 88 |
| 15 | Whitney | Anabelle | 83 | 15 | 98 |
| 16 | Wilma | Anabelle | 46 | 12 | 58 |
| 17 | Winnie | Anabelle | 37 | 10 | 47 |
| 18 | Wynona | Anabelle | 14 | 14 | 28 |
| 19 | Clarette | Cleopatra | 17 | 7 | 24 |
| 20 | Clarissa | Cleopatra | 21 | 6 | 27 |
| 21 | Claudine | Cleopatra | 12 | 5 | 17 |
| 22 | Clover | Cleopatra | 12 | 5 | 17 |
| 23 | Fiola | Fiola | 12 | 12 | 24 |
| 24 | Isabella | Fiola | 6 | 12 | 18 |
| 25 | Iseult | Fiola | 10 | 12 | 22 |
| 26 | Isis | Fiola | 11 | 15 | 26 |
| 27 | Ismena | Fiola | 18 | 5 | 23 |
| 28 | Arima | Kasturi | 24 | 16 | 40 |
| 29 | Kasturi | Kasturi | 274 | 96 | 370 |
| 30 | Kausalya | Kasturi | 196 | 102 | 298 |
| 31 | Keerthana | Kasturi | 108 | 98 | 206 |
| 32 | Ketki | Kasturi | 153 | 90 | 243 |
| 33 | Kokila | Kasturi | 89 | 85 | 174 |
| 34 | Koyna | Kasturi | 125 | 103 | 228 |
| 35 | Emerald | Katrina | 8 | 26 | 34 |
| 36 | Kaira | Katrina | 21 | 33 | 54 |
| 37 | Kate | Katrina | 6 | 4 | 10 |
| 38 | Meredith | Katrina | 14 | 8 | 22 |
| 39 | Merinda | Katrina | 7 | 36 | 43 |
| 40 | Merissa | Katrina | 16 | 30 | 46 |
| 41 | Lakshana | Lalanti | 9 | 10 | 19 |
| 42 | Lalanti | Lalanti | 5 | 7 | 12 |
| 43 | Lalasa | Lalanti | 9 | 5 | 14 |
| 44 | Jacintha | Lisa | 27 | 36 | 63 |
| 45 | Jackie | Lisa | 7 | 9 | 16 |
| 46 | Jeannie | Lisa | 13 | 21 | 34 |
| 47 | Joan | Lisa | 12 | 22 | 34 |
| 48 | Joycee | Lisa | 11 | 17 | 28 |
| 49 | Laika | Lisa | 54 | 43 | 97 |
| 50 | Laura | Lisa | 61 | 51 | 112 |
| 51 | Lauren | Lisa | 60 | 60 | 120 |
| 52 | Layla | Lisa | 16 | 6 | 22 |
| 53 | Leena | Lisa | 92 | 70 | 162 |
| 54 | Liezl | Lisa | 96 | 46 | 142 |
| 55 | Lily | Lisa | 166 | 41 | 207 |
| 56 | Linda | Lisa | 85 | 54 | 139 |
| 57 | Linette | Lisa | 30 | 21 | 51 |
| 58 | Lisa | Lisa | 74 | 13 | 87 |
| 59 | Lydia | Lisa | 96 | 79 | 175 |
| 60 | Lynn | Lisa | 66 | 77 | 143 |
| 61 | Ianthe | Menaka | 69 | 12 | 81 |
| 62 | Ilaena | Menaka | 41 | 21 | 62 |
| 63 | Iliora | Menaka | 65 | 21 | 86 |
| 64 | Ilsa | Menaka | 75 | 22 | 97 |
| 65 | Itzel | Menaka | 14 | 5 | 19 |
| 66 | Orianne | Menaka | 10 | 10 | 20 |
| 67 | Orlanda | Menaka | 7 | 10 | 17 |
| 68 | Midhuna | Mridula | 77 | 12 | 89 |
| 69 | Mihika | Mridula | 60 | 9 | 69 |
| 70 | Mihira | Mridula | 81 | 9 | 90 |
| 71 | Mitali | Mridula | 59 | 8 | 67 |
| 72 | Mithila | Mridula | 53 | 9 | 62 |
| 73 | Mridula | Mridula | 20 | 12 | 32 |
| 74 | Mridumita | Mridula | 51 | 12 | 63 |
| 75 | Ashley | Nakshatra | 24 | 13 | 37 |
| 76 | Gaargi | Nakshatra | 63 | 44 | 107 |
| 77 | Kai | Nakshatra | 34 | 22 | 56 |
| 78 | Kianna | Nakshatra | 35 | 14 | 49 |
| 79 | Nakshatra | Nakshatra | 71 | 56 | 127 |
| 80 | Namrata | Nakshatra | 59 | 45 | 104 |
| 81 | Nandini | Nakshatra | 59 | 50 | 109 |
| 82 | Narmada | Nakshatra | 37 | 42 | 79 |
| 83 | Navaneeta | Nakshatra | 20 | 19 | 39 |
| 84 | Naveena | Nakshatra | 36 | 25 | 61 |
| 85 | Navistha | Nakshatra | 10 | 6 | 16 |
| 86 | Navya | Nakshatra | 39 | 13 | 52 |
| 87 | Nayonika | Nakshatra | 21 | 18 | 39 |
| 88 | Nayra | Nakshatra | 20 | 6 | 26 |
| 89 | Anatolia | Olympia | 16 | 10 | 26 |
| 90 | Neena | Olympia | 14 | 6 | 20 |
| 91 | Raheema | Olympia | 19 | 10 | 29 |
| 92 | Rhoswen | Olympia | 8 | 5 | 13 |
| 93 | Rohini | Olympia | 11 | 11 | 22 |
| 94 | Ruhi | Olympia | 33 | 15 | 48 |
| 95 | Rumki | Olympia | 15 | 17 | 32 |
| 96 | Rupini | Olympia | 6 | 18 | 24 |
| 97 | SAF_with_Neena | Olympia | 16 | 6 | 22 |
| 98 | Zerlinda | Olympia | 7 | 5 | 12 |
| 99 | Amarilla | Osanna | 12 | 8 | 20 |
| 100 | Ambesafa | Osanna | 37 | 13 | 50 |
| 101 | Amelia | Osanna | 13 | 8 | 21 |
| 102 | Amy | Osanna | 20 | 7 | 27 |
| 103 | Deeksha | Osanna | 11 | 5 | 16 |
| 104 | Galina | Osanna | 33 | 15 | 48 |
| 105 | Galina_gp_SAF | Osanna | 12 | 5 | 17 |
| 106 | Garnet | Osanna | 26 | 6 | 32 |
| 107 | Gartred | Osanna | 5 | 8 | 13 |
| 108 | Gaynor | Osanna | 10 | 10 | 20 |
| 109 | Gina | Osanna | 10 | 14 | 24 |
| 110 | Osanna | Osanna | 28 | 6 | 34 |
| 111 | Camila | Patricia | 74 | 19 | 93 |
| 112 | Cancan | Patricia | 86 | 15 | 101 |
| 113 | Candice | Patricia | 121 | 20 | 141 |
| 114 | Cara | Patricia | 10 | 13 | 23 |
| 115 | Casey | Patricia | 72 | 50 | 122 |
| 116 | Cassandra | Patricia | 91 | 59 | 150 |
| 117 | Dana | Patricia | 224 | 28 | 252 |
| 118 | Dominique | Patricia | 165 | 48 | 213 |
| 119 | Doris | Patricia | 133 | 24 | 157 |
| 120 | Gemini | Patricia | 154 | 52 | 206 |
| 121 | Genette | Patricia | 163 | 67 | 230 |
| 122 | Genevieve | Patricia | 204 | 31 | 235 |
| 123 | Georgina | Patricia | 137 | 50 | 187 |
| 124 | Geraldine | Patricia | 36 | 6 | 42 |
| 125 | Gladys | Patricia | 83 | 20 | 103 |
| 126 | Glenda | Patricia | 57 | 7 | 64 |
| 127 | Gloria | Patricia | 32 | 6 | 38 |
| 128 | Pamela | Patricia | 166 | 47 | 213 |
| 129 | Patricia | Patricia | 158 | 42 | 200 |
| 130 | Paula | Patricia | 63 | 36 | 99 |
| 131 | Hannah | Tilottama | 41 | 55 | 96 |
| 132 | Marlene | Tilottama | 72 | 44 | 116 |
| 133 | Mary | Tilottama | 47 | 35 | 82 |
| 134 | Tilottama | Tilottama | 48 | 30 | 78 |
| 135 | Udbhasita | Unnati | 10 | 5 | 15 |
| 136 | Udyati | Unnati | 9 | 6 | 15 |
| 137 | Faiza | Victoria | 9 | 10 | 19 |
| 138 | Fawzia | Victoria | 30 | 19 | 49 |
| 139 | Floppy_ears | Victoria | 65 | 68 | 133 |
| 140 | Impedimenta | Victoria | 126 | 89 | 215 |
| 141 | Imperia | Victoria | 26 | 5 | 31 |
| 142 | Impetunia | Victoria | 90 | 76 | 166 |
| 143 | Ina | Victoria | 76 | 73 | 149 |
| 144 | Ipomoea | Victoria | 108 | 117 | 225 |
| 145 | Itamara | Victoria | 140 | 88 | 228 |
| 146 | Saina | Victoria | 25 | 14 | 39 |
| 147 | Sajida | Victoria | 133 | 72 | 205 |
| 148 | Salma | Victoria | 64 | 50 | 114 |
| 149 | Salvia | Victoria | 80 | 67 | 147 |
| 150 | Sameena | Victoria | 67 | 25 | 92 |
| 151 | Sarah | Victoria | 66 | 87 | 153 |
| 152 | Sarayu | Victoria | 62 | 72 | 134 |
| 153 | Sarin | Victoria | 62 | 52 | 114 |
| 154 | Saroj | Victoria | 41 | 33 | 74 |
| 155 | Serena | Victoria | 69 | 85 | 154 |
| 156 | Sucheta | Victoria | 30 | 24 | 54 |
| 157 | Sunaina | Victoria | 85 | 67 | 152 |
| 158 | Suneeti | Victoria | 61 | 48 | 109 |
| 159 | Sunetra | Victoria | 87 | 106 | 193 |
| 160 | Supri | Victoria | 75 | 53 | 128 |
| 161 | Surabhi | Victoria | 29 | 25 | 54 |
| 162 | Sushma | Victoria | 47 | 63 | 110 |
| 163 | Sushmita | Victoria | 31 | 19 | 50 |
| 164 | Suvarti | Victoria | 90 | 88 | 178 |
| 165 | Suveera | Victoria | 86 | 169 | 255 |
| 166 | Suvrata | Victoria | 82 | 110 | 192 |
| 167 | Ursula | Victoria | 31 | 66 | 97 |
| 168 | Valerie | Victoria | 125 | 134 | 259 |
| 169 | Vanessa | Victoria | 192 | 136 | 328 |
| 170 | Vera | Victoria | 31 | 36 | 67 |
| 171 | Victoria | Victoria | 21 | 26 | 47 |

Supplementary Material S3

*Calf present and calf absent periods within each clan*

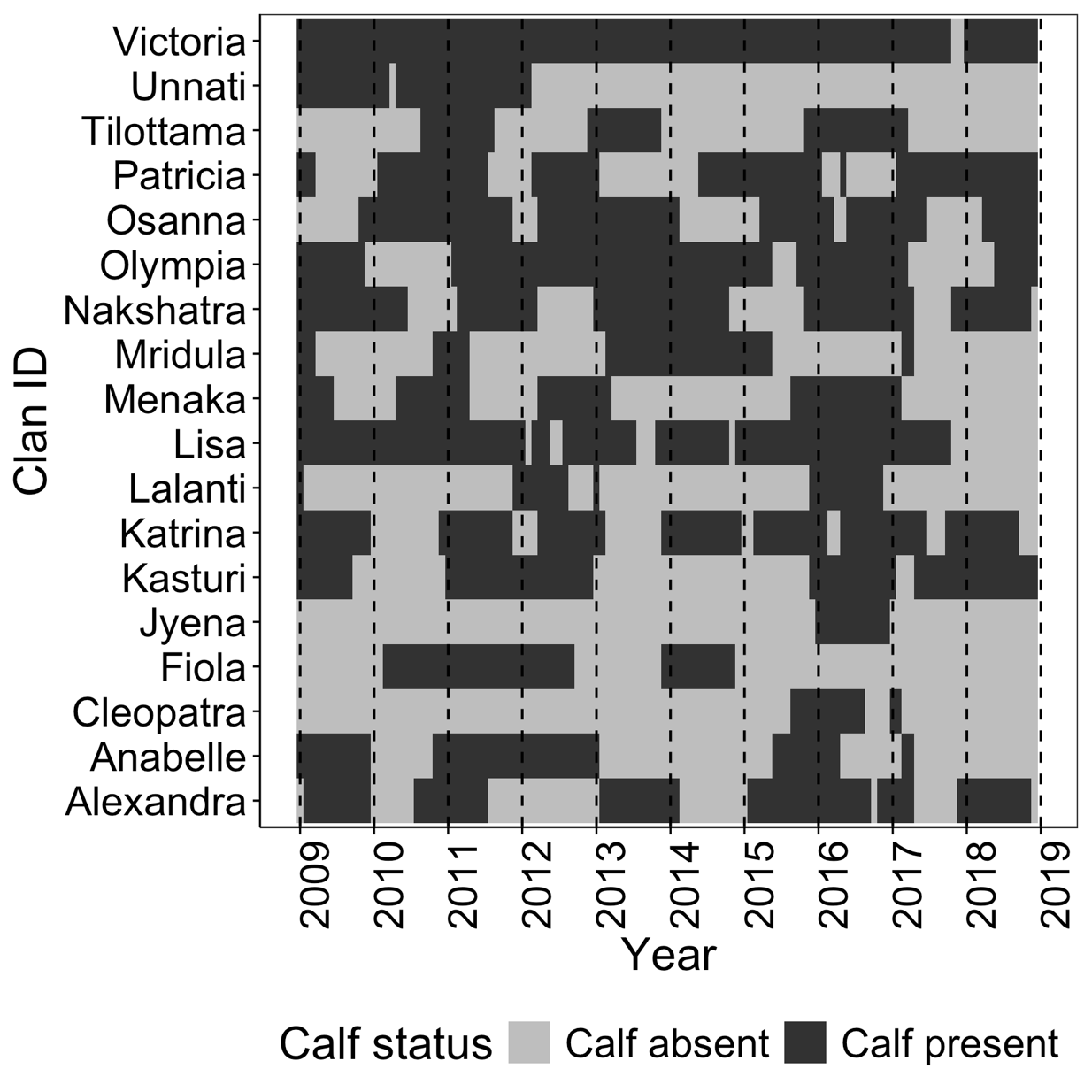

**Supplementary Material S3, Figure S1**. Periods of calf presence and absence across all the sampled clans during the entire study period. Most clans did not have a calf-present and a calf-absent period within each sampling year. Additionally, females were sighted opportunistically. Therefore, it was difficult to accumulate five sightings in both calf presence and absence within each sampling year for each female. As a result, we conducted the association analysis over the entire sampling period. Furthermore, note that the objective of this study was to test the overall effect of calf absence/presence on female social structure rather than testing the year-to-year stability in the effect of this predictor (see Methods). We plan to do the stability analysis when we accumulate sufficient data.

Supplementary Material S4

*Group size model structures and results*

**Supplementary Material S4, Table S1**. Model structure and results of m_AF, i.e., to predict the number of adult females in a group (the dependent variable). Estimated mean, standard error (SE), lower and upper 95% CI, and *Z* and *P* values are provided for the fixed predictors in log link space. The number of groups and the estimated variance and standard deviation (SD) of the random intercepts and slopes are provided for each grouping factor. Calf status has two levels (absence/presence) with calf absence as the reference level. (P) refers to calf presence. (1 + Calf status || Clan ID) allows for different clans to have different baseline levels (random intercept) of the dependent variable, as well as for different clans to have different effects of calf status (random slope) on the dependent variable. || means that the correlation parameter between the random intercept and the random slope of the grouping factor was not estimated. (1 + Calf status + Number of adult females in the clan in the sampling year* || Year) allows for baseline levels of the dependent variable to vary among years (random intercept), as well as for the effect of calf status and the number of adult females in the clan on the dependent variable to vary among years (random slopes). We used a negative binomial error distribution for the dependent variable. *z-transformed to a mean of 0 and a SD of 1. *P* values were obtained using the *drop1* function. Significant *P* values are shown in bold. ^†^Not shown due to limited interpretability.

| ***Number of adult females in a group (R^2^m: 0.218; R^2^c: 0.339)*** | | | | | | |
| --- | --- | --- | --- | --- | --- | --- |
| m_AF: glmmTMB(Number of adult females in a group ~ Calf status + Number of adult females in the clan in the sampling year* + (1 + Calf status \|\| Clan ID) + (1 + Calf status + Number of adult females in the clan in the sampling year* \|\| Year), family = "nbinom2") | | | | | | |
| **Variable** | **Estimate** | **Estimated**  **SE** | **Lower 95% CI** | **Upper 95% CI** | ***Z*** | ***P*** |
| Intercept | 0.770 | 0.047 | 0.678 | 0.862 | † | † |
| Calf status (P) | 0.673 | 0.094 | 0.488 | 0.857 | 7.151 | **<0.001** |
| Number of adult females in the clan* | 0.113 | 0.047 | 0.022 | 0.204 | 2.434 | **0.036** |
| **Grouping factor** | **Random intercept** | **Random slope** | **Estimated variance** | **Estimated SD** | **Estimated correlation** | **Number of groups** |
| Clan ID | Intercept | _ | 0.000 | 0.000 | _ | 14 clans |
|  | _ | Calf status (P) | 0.047 | 0.218 | _ | _ |
| Year | Intercept | _ | 0.019 | 0.136 | _ | 10 years |
|  | _ | Calf status (P) | 0.039 | 0.199 | _ | _ |
|  | _ | Number of adult females in the clan* | 0.016 | 0.127 | _ | _ |

The original mean (SD) of the number of adult females in the clan was 20.29 (8.591).

**Supplementary Material S4, Table S2**. Model structure and results of m_YAF, i.e., to predict the number of young adult females in a group (the dependent variable). Estimated mean, standard error (SE), lower and upper 95% CI, and *Z* and *P* values are provided for the fixed predictors in log link space. The number of groups and the estimated variance and standard deviation (SD) of the random intercepts and slopes are provided for each grouping factor. Calf status has two levels (absence/presence) with calf absence as the reference level. (P) refers to calf presence. (1 + Calf status || Clan ID) allows for different clans to have different baseline levels (random intercept) of the dependent variable, as well as for different clans to have different effects of calf status (random slope) on the dependent variable. || means that the correlation parameter between the random intercept and the random slope of the grouping factor was not estimated. (1 + Calf status + Number of young adult females in the clan in the sampling year* || Year) allows for baseline levels of the dependent variable to vary among years (random intercept), as well as for the effect of calf status and the number of young adult females in the clan on the dependent variable to vary among years (random slopes). ^†^Not shown due to limited interpretability. *z-transformed to a mean of 0 and a SD of 1. *P* values were obtained using the *drop1* function. Significant *P* values are shown in bold.

| ***Number of young adult females in a group (R^2^m: 0.400; R^2^c: 0.593)*** | | | | | | |
| --- | --- | --- | --- | --- | --- | --- |
| m_YAF: glmmTMB(Number of young adult females in a group ~ Calf status + Number of young adult females in the clan in the sampling year* + (1 + Calf status \|\| Clan ID) + (1 + Calf status + Number of young adult females in the clan in the sampling year* \|\| Year), family = "poisson") | | | | | | |
| **Variable** | **Estimate** | **Estimated** | **Lower 95% CI** | **Upper 95% CI** | ***Z*** | ***P*** |
|  |  | **SE** |  |  |  |  |
| Intercept | -0.732 | 0.163 | -1.052 | -0.412 | † | † |
| Calf status (P) | 0.791 | 0.161 | 0.475 | 1.107 | 4.907 | **<0.001** |
| Number of young adult females in the clan* | 0.793 | 0.136 | 0.526 | 1.060 | 5.821 | **<0.001** |
| **Grouping factor** | **Random intercept** | **Random slope** | **Estimated variance** | **Estimated SD** | **Estimated correlation** | **Number of groups** |
| Clan ID | Intercept | _ | 0.190 | 0.436 | _ | 14 clans |
|  | _ | Calf status (P) | 0.167 | 0.408 | _ | _ |
| Year | Intercept | _ | 0.053 | 0.230 | _ | 10 years |
|  | _ | Calf status (P) | 0.061 | 0.248 | _ | _ |
|  | _ | Number of young adult females in the clan* | 0.068 | 0.260 | _ | _ |

The original mean (SD) of the number of young adult females in the clan was 4.80 (3.274).

**Supplementary Material S4, Table S3**. Model structure and results of m_SAF, i.e., to predict the number of subadult females in a group (the dependent variable). Estimated mean, standard error (SE), lower and upper 95% CI, and *Z* and *P* values are provided for the fixed predictors in log link space. The number of groups and the estimated variance and standard deviation (SD) of the random intercepts and slopes are provided for each grouping factor. Calf status has two levels (absence/presence) with calf absence as the reference level. (P) refers to calf presence. (1 + Calf status || Clan ID) allows for different clans to have different baseline levels (random intercept) of the dependent variable, as well as for different clans to have different effects of calf status (random slope) on the dependent variable. || means that the correlation parameter between the random intercept and the random slope of the grouping factor was not estimated. (1 + Calf status + Number of subadult females in the clan in the sampling year* || Year) allows for baseline levels of the dependent variable to vary among years (random intercept), as well as for the effect of calf status and the number of subadult females in the clan on the dependent variable to vary among years (random slopes). ^†^Not shown due to limited interpretability. *z-transformed to a mean of 0 and a SD of 1. *P* values were obtained using the *drop1* function. Significant *P* values are shown in bold.

| ***Number of subadult females in a group (R^2^m: 0.341; R^2^c: 0.723)*** | | | | | | |
| --- | --- | --- | --- | --- | --- | --- |
| m_YAF: glmmTMB(Number of subadult females in a group ~ Calf status + Number of subadult females in the clan in the sampling year* + (1 + Calf status \|\| Clan ID) + (1 + Calf status + Number of subadult females in the clan in the sampling year* \|\| Year), family = "poisson") | | | | | | |
| **Variable** | **Estimate** | **Estimated** | **Lower 95% CI** | **Upper 95% CI** | ***Z*** | ***P*** |
|  |  | **SE** |  |  |  |  |
| Intercept | -0.974 | 0.213 | -1.392 | -0.556 | † | † |
| Calf status (P) | 0.924 | 0.159 | 0.613 | 1.236 | 5.813 | **<0.001** |
| Number of subadult females in the clan* | 0.870 | 0.313 | 0.256 | 1.483 | 2.778 | **0.012** |
| **Grouping factor** | **Random intercept** | **Random slope** | **Estimated variance** | **Estimated SD** | **Estimated correlation** | **Number of groups** |
| Clan ID | Intercept | _ | 0.254 | 0.504 | _ | 14 clans |
|  | _ | Calf status (P) | 0.184 | 0.428 | _ | _ |
| Year | Intercept | _ | 0.096 | 0.310 | _ | 10 years |
|  | _ | Calf status (P) | 0.029 | 0.171 | _ | _ |
|  | _ | Number of subadult females in the clan* | 0.678 | 0.824 | _ | _ |

The original mean (SD) of the number of subadult females in the clan was 5.09 (5.595).

**Supplementary Material S4, Table S4**. Model structure and results of m_AF1, i.e., to predict the number of adult females in a group without their own dependent offspring (the dependent variable). Estimated mean, standard error (SE), lower and upper 95% CI, and *Z* and *P* values are provided for the fixed predictors in log link space. The number of groups and the estimated variance and standard deviation (SD) of the random intercepts and slopes are provided for each grouping factor. Calf status has two levels (absence/presence) with calf absence as the reference level. (P) refers to calf presence. (1 + Calf status || Clan ID) allows for different clans to have different baseline levels (random intercept) of the dependent variable, as well as for different clans to have different effects of calf status (random slope) on the dependent variable. || means that the correlation parameter between the random intercept and the random slope of the grouping factor was not estimated. (1 + Calf status + Number of adult females in the clan in the sampling year* || Year) allows for baseline levels of the dependent variable to vary among years (random intercept), as well as for the effect of calf status and the number of adult females in the clan on the dependent variable to vary among years (random slopes). ^†^Not shown due to limited interpretability. *z-transformed to a mean of 0 and a SD of 1. *P* values were obtained using the *drop1* function. Significant *P* values are shown in bold.

| ***Number of adult females in a group without their own dependent offspring***  ***(R^2^m: 0.069; R^2^c: 0.222)*** | | | | | | |
| --- | --- | --- | --- | --- | --- | --- |
| m_AF1: glmmTMB(Number of adult females in a group without their own dependent offspring ~ Calf status + Number of adult females in the clan in the sampling year* + (1 + Calf status \|\| Clan ID) + (1 + Calf status + Number of adult females in the clan in the sampling year* \|\| Year), family = "nbinom2") | | | | | | |
| **Variable** | **Estimate** | **Estimated** | **Lower 95% CI** | **Upper 95% CI** | ***Z*** | ***P*** |
|  |  | **SE** |  |  |  |  |
| Intercept | 0.666 | 0.043 | 0.582 | 0.751 | † | † |
| Calf status (P) | 0.356 | 0.127 | 0.108 | 0.604 | 2.814 | **0.009** |
| Number of adult females in the clan* | 0.106 | 0.057 | -0.006 | 0.218 | 1.862 | 0.084 |
| **Grouping factor** | **Random intercept** | **Random slope** | **Estimated variance** | **Estimated SD** | **Estimated correlation** | **Number of groups** |
| Clan ID | Intercept | _ | 0.002 | 0.048 | _ | 14 clans |
|  | _ | Calf status (P) | 0.087 | 0.295 | _ | _ |
| Year | Intercept | _ | 0.009 | 0.096 | _ | 10 years |
|  | _ | Calf status (P) | 0.074 | 0.272 | _ | _ |
|  | _ | Number of adult females in the clan* | 0.022 | 0.149 | _ | _ |

The original mean (SD) of the number of adult females in the clan was 20.29 (8.591).

**Supplementary Material S4, Table S5**. Model structure and results of m_YAF1, i.e., to predict the number of young adult females in a group without their own dependent offspring (the dependent variable). Estimated mean, standard error (SE), lower and upper 95% CI, and *Z* and *P* values are provided for the fixed predictors in log link space. The number of groups and the estimated variance and standard deviation (SD) of the random intercepts and slopes are provided for each grouping factor. Calf status has two levels (absence/presence) with calf absence as the reference level. (P) refers to calf presence. (1 + Calf status || Clan ID) allows for different clans to have different baseline levels (random intercept) of the dependent variable, as well as for different clans to have different effects of calf status (random slope) on the dependent variable. || means that the correlation parameter between the random intercept and the random slope of the grouping factor was not estimated. (1 + Calf status + Number of young adult females in the clan in the sampling year* || Year) allows for baseline levels of the dependent variable to vary among years (random intercept), as well as for the effect of calf status and the number of young adult females in the clan on the dependent variable to vary among years (random slopes). ^†^Not shown due to limited interpretability. *z-transformed to a mean of 0 and a SD of 1. *P* values were obtained using the *drop1* function. Significant *P* values are shown in bold.

| ***Number of young adult females in a group without their own dependent offspring***  ***(R^2^m: 0.371; R^2^c: 0.611)*** | | | | | | |
| --- | --- | --- | --- | --- | --- | --- |
| m_YAF1: glmmTMB(Number of young adult females in a group without their own dependent offspring ~ Calf status + Number of young adult females in the clan in the sampling year* + (1 + Calf status \|\| Clan ID) + (1 + Calf status + Number of young adult females in the clan in the sampling year* \|\| Year), family = "poisson") | | | | | | |
| **Variable** | **Estimate** | **Estimated** | **Lower 95% CI** | **Upper 95% CI** | ***Z*** | ***P*** |
|  |  | **SE** |  |  |  |  |
| Intercept | -0.754 | 0.185 | -1.117 | -0.391 | † | † |
| Calf status (P) | 0.612 | 0.199 | 0.221 | 1.002 | 3.069 | **0.008** |
| Number of young adult females in the clan* | 0.910 | 0.175 | 0.567 | 1.253 | 5.205 | **<0.001** |
| **Grouping factor** | **Random intercept** | **Random slope** | **Estimated variance** | **Estimated SD** | **Estimated correlation** | **Number of groups** |
| Clan ID | Intercept | _ | 0.241 | 0.491 | _ | 14 clans |
|  | _ | Calf status (P) | 0.355 | 0.596 | _ | _ |
| Year | Intercept | _ | 0.078 | 0.279 | _ | 10 years |
|  | _ | Calf status (P) | 0.044 | 0.210 | _ | _ |
|  | _ | Number of young adult females in the clan* | 0.154 | 0.392 | _ | _ |

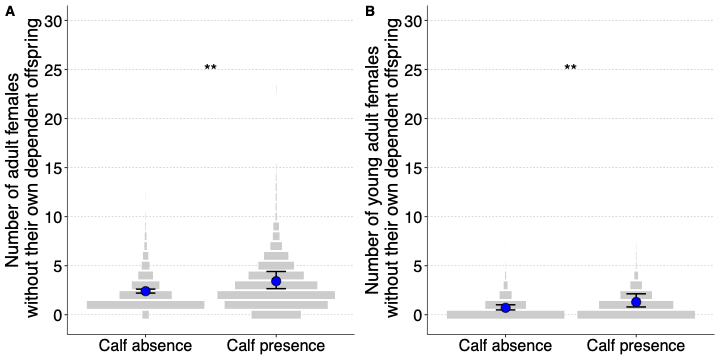

**Supplementary Material S4, Figure S1**. Effect of calves on female group size. Mean model predictions (blue dot) and the confidence intervals (black bars) are shown for the numbers of (A) adult and (B) young adult females in a group without their own dependent offspring. The observed data are as shown as discrete violin plots (grey bars). The widths of the bars show the relative frequency of observations within each calf status, and width of the widest bar is scaled to be the same between calf presence and absence. Significance codes: <0.001 ‘***’ 0.001 ‘**’ 0.01 ‘*’ 0.05 ‘.’.

Supplementary Material S5

*Experienced group size model structure and results*

**Supplementary Material S5, Table S1**. Model structure and results of mE_AF. The dependent variable represents the number of adult females experienced by adult females in groups with and without calves. Estimated mean, standard error (SE), lower and upper 95% CI, and *Z* and *P* values are provided for the fixed predictors in log link space. The number of groups and the estimated variance and standard deviation (SD) of the random intercepts and slopes are provided for each grouping factor. Calf status has two levels (absence/presence) with calf absence as the reference level. (P) refers to calf presence. (1 + Calf status || Adult female ID) allows for different females to have different baseline levels (random intercept) of the dependent variable, as well as for different females to have different effects of calf status (random slope) on the dependent variable. || means that the correlation parameter between the random intercept and the random slope of the grouping factor was not estimated. (1 + Calf status + Number of adult females in the clan in the sampling year* || Year) allows for baseline levels of the dependent variable to vary among years (random intercept), as well as for the effect of calf status and the number of adult females in the clan on the dependent variable to vary among years (random slopes). ^†^Not shown due to limited interpretability. *z-transformed to a mean of 0 and a SD of 1. *P* values were obtained using the *drop1* function. Significant *P* values are shown in bold.

| ***Experienced number of adult females (R^2^m: 0.202; R^2^c: 0.713)*** | | | | | | |
| --- | --- | --- | --- | --- | --- | --- |
| mE_AF: glmmTMB(Experienced number of adult females ~ Calf status + Number of adult females in the clan in the sampling year* + (1 + Calf status \|\| Adult female ID) + (1 + Calf status \|\| Clan ID) + (1 + Calf status + Number of adult females in the clan in the sampling year* \|\| Year) + (1 \| Sighting ID), family = "truncated_poisson") | | | | | | |
| **Variable** | **Estimate** | **Estimated** | **Lower 95% CI** | **Upper 95% CI** | ***Z*** | ***P*** |
|  |  | **SE** |  |  |  |  |
| Intercept | 0.815 | 0.126 | 0.569 | 1.061 | † | † |
| Calf status (P) | 0.711 | 0.122 | 0.472 | 0.951 | 5.827 | **<0.001** |
| Number of adult females in the clan* | 0.166 | 0.114 | -0.056 | 0.389 | 1.465 | 0.140 |
| **Grouping factor** | **Random intercept** | **Random**  **slope** | **Estimated variance** | **Estimated SD** | **Estimated correlation** | **Number of groups** |
| Adult female ID | Intercept | _ | 0.001 | 0.025 | _ | 84 adult females |
|  |  | Calf status (P) | 0.000 | 0.000 |  |  |
| Clan ID | Intercept | _ | 0.031 | 0.175 | _ | 6 clans |
|  | _ | Calf status (P) | 0.045 | 0.212 | _ | _ |
| Year | Intercept | _ | 0.051 | 0.225 | _ | 9 years |
|  | _ | Calf status (P) | 0.045 | 0.212 | _ | _ |
|  | _ | Number of adult females in the clan* | 0.035 | 0.187 | _ | _ |
| Sighting ID | Intercept | _ | 0.228 | 0.477 | _ | 1946 sightings |

The original mean (SD) of the number of adult females in the clan was 22.23 (9.662).

**Supplementary Material S5, Table S2**. Model structure and results of mE_YAF. The dependent variable represents the number of young adult females experienced by adult females in groups with and without calves. Estimated mean, standard error (SE), lower and upper 95% CI, and *Z* and *P* values are provided for the fixed predictors in log link space. The number of groups and the estimated variance and standard deviation (SD) of the random intercepts and slopes are provided for each grouping factor. Calf status has two levels (absence/presence) with calf absence as the reference level. (P) refers to calf presence. (1 + Calf status || Adult female ID) allows for different females to have different baseline levels (random intercept) of the dependent variable, as well as for different females to have different effects of calf status (random slope) on the dependent variable. || means that the correlation parameter between the random intercept and the random slope of the grouping factor was not estimated. (1 + Calf status + Number of young adult females in the clan in the sampling year* || Year) allows for baseline levels of the dependent variable to vary among years (random intercept), as well as for the effect of calf status and the number of young adult females in the clan on the dependent variable to vary among years (random slopes). ^†^Not shown due to limited interpretability. *z-transformed to a mean of 0 and a SD of 1. *P* values were obtained using the *drop1* function. Significant *P* values are shown in bold.

| ***Experienced number of young adult females (R^2^m: 0.239; R^2^c: 0.682)*** | | | | | | |
| --- | --- | --- | --- | --- | --- | --- |
| mE_YAF: glmmTMB(Experienced number of young adult females ~ Calf status + Number of young adult females in the clan in the sampling year* + (1 + Calf status \|\| Adult female ID) + (1 + Calf status \|\| Clan ID) + (1 + Calf status + Number of young adult females in the clan in the sampling year* \|\| Year) + (1 \| Sighting ID), family = "poisson") | | | | | | |
| **Variable** | **Estimate** | **Estimated** | **Lower 95% CI** | **Upper 95% CI** | ***Z*** | ***P*** |
|  |  | **SE** |  |  |  |  |
| Intercept | -0.674 | 0.233 | -1.130 | -0.217 | † | † |
| Calf status (P) | 0.617 | 0.203 | 0.218 | 1.015 | 3.033 | **0.025** |
| Number of young adult females in the clan* | 0.613 | 0.185 | 0.250 | 0.975 | 3.312 | **0.003** |
| **Grouping factor** | **Random intercept** | **Random slope** | **Estimated variance** | **Estimated SD** | **Estimated correlation** | **Number of groups** |
| Adult female ID | Intercept | _ | 0.028 | 0.166 | _ | 84 adult females |
|  |  | Calf status (P) | 0.000 | 0.000 |  |  |
| Clan ID | Intercept | _ | 0.058 | 0.241 | _ | 6 clans |
|  | _ | Calf status (P) | 0.123 | 0.351 | _ | _ |
| Year | Intercept | _ | 0.290 | 0.538 | _ | 9 years |
|  | _ | Calf status (P) | 0.096 | 0.310 | _ | _ |
|  | _ | Number of young adult females in the clan* | 0.086 | 0.294 | _ | _ |
| Sighting ID | Intercept | _ | 0.333 | 0.577 | _ | 1946 sightings |

The original mean (SD) of the number of young adult females in the clan was 5.45 (3.633).

**Supplementary Material S5, Table S3**. Model structure and results of mE_SAF. The dependent variable represents the number of subadult females experienced by adult females in groups with and without calves. Estimated mean, standard error (SE), lower and upper 95% CI, and *Z* and *P* values are provided for the fixed predictors in log link space. The number of groups and the estimated variance and standard deviation (SD) of the random intercepts and slopes are provided for each grouping factor. Calf status has two levels (absence/presence) with calf absence as the reference level. (P) refers to calf presence. (1 + Calf status || Adult female ID) allows for different females to have different baseline levels (random intercept) of the dependent variable, as well as for different females to have different effects of calf status (random slope) on the dependent variable. || means that the correlation parameter between the random intercept and the random slope of the grouping factor was not estimated. (1 + Calf status + Number of subadult females in the clan in the sampling year* || Year) allows for baseline levels of the dependent variable to vary among years (random intercept), as well as for the effect of calf status and the number of subadult females in the clan on the dependent variable to vary among years (random slopes). ^†^Not shown due to limited interpretability. *z-transformed to a mean of 0 and a SD of 1. *P* values were obtained using the *drop1* function. Significant *P* values are shown in bold.

| ***Experienced number of subadult females (R^2^m: 0.042; R^2^c: 0.945)*** | | | | | | |
| --- | --- | --- | --- | --- | --- | --- |
| mE_SAF: glmmTMB(Experienced number of subadult females ~ Calf status + Number of subadult females in the clan in the sampling year* + (1 + Calf status \|\| Adult female ID) + (1 + Calf status \|\| Clan ID) + (1 + Calf status + Number of subadult females in the clan in the sampling year* \|\| Year) + (1 \| Sighting ID), family = "poisson") | | | | | | |
| **Variable** | **Estimate** | **Estimated** | **Lower 95% CI** | **Upper 95% CI** | ***Z*** | ***P*** |
|  |  | **SE** |  |  |  |  |
| Intercept | -1.229 | 0.687 | -2.575 | 0.118 | † | † |
| Calf status (P) | 0.846 | 0.245 | 0.366 | 1.325 | 3.457 | **0.007** |
| Number of subadult females in the clan* | 0.310 | 0.833 | -1.323 | 1.943 | 0.372 | 0.709 |
| **Grouping factor** | **Random intercept** | **Random slope** | **Estimated variance** | **Estimated SD** | **Estimated correlation** | **Number of groups** |
| Adult female ID | Intercept | _ | 0.022 | 0.147 | _ | 84 adult females |
|  |  | Calf status (P) | 0.000 | 0.000 |  |  |
| Clan ID | Intercept | _ | 1.760 | 1.327 | _ | 6 clans |
|  | _ | Calf status (P) | 0.220 | 0.469 | _ | _ |
| Year | Intercept | _ | 0.453 | 0.673 | _ | 9 years |
|  | _ | Calf status (P) | 0.052 | 0.229 | _ | _ |
|  | _ | Number of subadult females in the clan* | 3.437 | 1.854 | _ | _ |
| Sighting ID | Intercept | _ | 0.341 | 0.584 | _ | 1946 sightings |

The original mean (SD) of the number of subadult females in the clan was 7.02 (6.306).

**Supplementary Material S5, Table S4**. Model structure and results of mE_AF1. The dependent variable represents the number of adult females without their own dependent offspring experienced by adult females in groups with and without calves. Estimated mean, standard error (SE), lower and upper 95% CI, and *Z* and *P* values are provided for the fixed predictors in log link space. The number of groups and the estimated variance and standard deviation (SD) of the random intercepts and slopes are provided for each grouping factor. Calf status has two levels (absence/presence) with calf absence as the reference level. (P) refers to calf presence. (1 + Calf status || Adult female ID) allows for different females to have different baseline levels (random intercept) of the dependent variable, as well as for different females to have different effects of calf status (random slope) on the dependent variable. || means that the correlation parameter between the random intercept and the random slope of the grouping factor was not estimated. (1 + Calf status + Number of adult females in the clan in the sampling year* || Year) allows for baseline levels of the dependent variable to vary among years (random intercept), as well as for the effect of calf status and the number of adult females in the clan on the dependent variable to vary among years (random slopes). ^†^Not shown due to limited interpretability. *z-transformed to a mean of 0 and a SD of 1. *P* values were obtained using the *drop1* function. Significant *P* values are shown in bold.

| ***Experienced number of adult females without their own dependent offspring***  ***(R^2^m: 0.130; R^2^c: 0.751)*** | | | | | | |
| --- | --- | --- | --- | --- | --- | --- |
| mE_AF1: glmmTMB(Experienced number of adult females without their own dependent offspring ~ Calf status + Number of adult females in the clan in the sampling year* + (1 + Calf status \|\| Adult female ID) + (1 + Calf status \|\| Clan ID) + (1 + Calf status + Number of adult females in the clan in the sampling year* \|\| Year) + (1 \| Sighting ID), family = "poisson") | | | | | | |
| **Variable** | **Estimate** | **Estimated** | **Lower 95% CI** | **Upper 95% CI** | ***Z*** | ***P*** |
|  |  | **SE** |  |  |  |  |
| Intercept | 0.283 | 0.212 | -0.133 | 0.700 | † | † |
| Calf status (P) | 0.637 | 0.131 | 0.379 | 0.894 | 4.849 | **0.001** |
| Number of adult females in the clan* | 0.240 | 0.199 | -0.150 | 0.630 | 1.207 | 0.188 |
| **Grouping factor** | **Random intercept** | **Random slope** | **Estimated variance** | **Estimated SD** | **Estimated correlation** | **Number of groups** |
| Adult female ID | Intercept | _ | 0.013 | 0.115 | _ | 84 adult females |
|  |  | Calf status (P) | 0.000 | 0.000 |  |  |
| Clan ID | Intercept | _ | 0.093 | 0.306 | _ | 6 clans |
|  | _ | Calf status (P) | 0.043 | 0.206 | _ | _ |
| Year | Intercept | _ | 0.124 | 0.352 | _ | 9 years |
|  | _ | Calf status (P) | 0.057 | 0.240 | _ | _ |
|  | _ | Number of adult females in the clan* | 0.076 | 0.276 | _ | _ |
| Sighting ID | Intercept | _ | 0.416 | 0.645 | _ | 1946 sightings |

The original mean (SD) of the number of adult females in the clan was 22.23 (9.662).

**Supplementary Material S5, Table S5**. Model structure and results of mE_YAF1. The dependent variable represents the number of young adult females without their own dependent offspring experienced by adult females in groups with and without calves. Estimated mean, standard error (SE), lower and upper 95% CI, and *Z* and *P* values are provided for the fixed predictors in log link space. The number of groups and the estimated variance and standard deviation (SD) of the random intercepts and slopes are provided for each grouping factor. Calf status has two levels (absence/presence) with calf absence as the reference level. (P) refers to calf presence. (1 + Calf status || Adult female ID) allows for different females to have different baseline levels (random intercept) of the dependent variable, as well as for different females to have different effects of calf status (random slope) on the dependent variable. || means that the correlation parameter between the random intercept and the random slope of the grouping factor was not estimated. (1 + Calf status + Number of young adult females in the clan in the sampling year* || Year) allows for baseline levels of the dependent variable to vary among years (random intercept), as well as for the effect of calf status and the number of young adult females in the clan on the dependent variable to vary among years (random slopes). ^†^Not shown due to limited interpretability. *z-transformed to a mean of 0 and a SD of 1. *P* values were obtained using the *drop1* function. Significant *P* values are shown in bold.

| ***Experienced number of young adult females without their own dependent offspring***  ***(R^2^m: 0.417; R^2^c: 0.790)*** | | | | | | |
| --- | --- | --- | --- | --- | --- | --- |
| mE_YAF1: glmmTMB(Experienced number of young adult females without their own dependent offspring ~ Calf status + Number of young adult females in the clan in the sampling year* + (1 + Calf status \|\| Adult female ID) + (1 + Calf status \|\| Clan ID) + (1 + Calf status + Number of young adult females in the clan in the sampling year* \|\| Year) + (1 \| Sighting ID), family = "poisson") | | | | | | |
| **Variable** | **Estimate** | **Estimated** | **Lower 95% CI** | **Upper 95% CI** | ***Z*** | ***P*** |
|  |  | **SE** |  |  |  |  |
| Intercept | -1.160 | 0.294 | -1.736 | -0.584 | † | † |
| Calf status (P) | 0.679 | 0.265 | 0.160 | 1.198 | 2.563 | **0.039** |
| Number of young adult females in the clan* | 1.345 | 0.432 | 0.498 | 2.193 | 3.113 | **0.002** |
| **Grouping factor** | **Random intercept** | **Random slope** | **Estimated variance** | **Estimated SD** | **Estimated correlation** | **Number of groups** |
| Adult female ID | Intercept | _ | 0.092 | 0.304 | _ | 84 adult females |
|  |  | Calf status (P) | 0.000 | 0.000 |  |  |
| Clan ID | Intercept | _ | 0.188 | 0.434 | _ | 6 clans |
|  | _ | Calf status (P) | 0.160 | 0.399 | _ | _ |
| Year | Intercept | _ | 0.067 | 0.259 | _ | 9 years |
|  | _ | Calf status (P) | 0.242 | 0.492 | _ | _ |
|  | _ | Number of young adult females in the clan* | 0.665 | 0.816 | _ | _ |
| Sighting ID | Intercept | _ | 0.541 | 0.736 | _ | 1946 sightings |

The original mean (SD) of the number of young adult females in the clan was 5.45 (3.633).

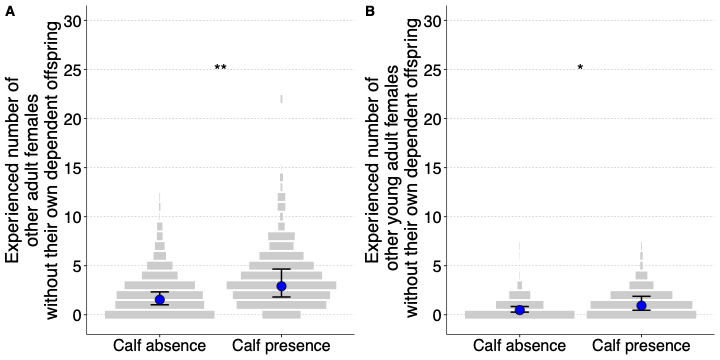

**Supplementary Material S5, Figure S1.** Effect of calves on female experienced group size. Mean model predictions (blue dot) and the confidence intervals (black bars) are shown for the experienced numbers of (A) adult and (B) young adult females without their own dependent offspring. The observed data are as shown as discrete violin plots (grey bars). The widths of the bars show the relative frequency of observations within each calf status, and width of the widest bar is scaled to be the same between calf presence and absence. Significance codes: <0.001 ‘***’ 0.001 ‘**’ 0.01 ‘*’ 0.05 ‘.’.

Supplementary Material S6

*Female association in calf presence and absence*

**Supplementary Material S6, Table S1**. Average (SD) AI, kurtosis of AI, and network statistics in the presence and absence of calves, and averages of these statistics from sampled randomisation tests are provided. The analyses were carried out using sightings of 172 unique adult females. Significant *P* values are shown in bold. The permuted SD values are average SD values (averaged across 10,000 randomised datasets).

| **Category** | **Average AI (SD)** | **Kurtosis of AI** | **Average degree (SD)** | **Average clustering coefficient (SD)** |
| --- | --- | --- | --- | --- |
| Calf absence observed | 0.007 (0.0425) | 108.07 | 11.9 (9.89) | 0.91 (0.117) |
| Calf presence observed | 0.013 (0.0694) | 71.68 | 12.5 (9.91) | 0.91 (0.125) |
| Calf absence permuted | 0.009 (0.0496) | 87.13 | 13.2 (10.28) | 0.93 (0.102) |
| Calf presence permuted | 0.009 (0.0514) | 94.07 | 11.3 (9.29) | 0.89 (0.130) |
| *P* value | **<0.001** | **<0.001** | **<0.001** | 0.997 |
